# Temperature explains broad patterns of Ross River virus transmission

**DOI:** 10.1101/286724

**Authors:** Marta S. Shocket, Sadie J. Ryan, Erin A. Mordecai

## Abstract

Thermal biology predicts that vector-borne disease transmission peaks at intermediate temperatures and declines at high and low temperatures. However, thermal optima and limits remain unknown for most vector-borne pathogens. We built a mechanistic model for the thermal response of Ross River virus, an important mosquito-borne pathogen in Australia, Pacific Islands, and potentially emerging worldwide. Transmission peaks at moderate temperatures (26.4°C) and declines to zero at thermal limits (17.0°C and 31.5°C). The model accurately predicts that transmission is year-round endemic in the tropics but seasonal in temperate areas, and matches the nationwide seasonal peak in human cases. Climate warming will likely increase transmission in temperate areas (where most Australians live) but decrease transmission in tropical areas where mean temperatures are already near the thermal optimum. These results illustrate the importance of nonlinear models for inferring the role of temperature in disease dynamics and predicting responses to climate change.

**Authorship statement:** EAM conceived of and designed the study. MSS collected data, performed statistical analyses, and wrote the first draft of the manuscript. SJR performed geographic analyses. All authors revised and approved the manuscript.

**Data accessibility statement:** Upon acceptance, all data will be archived in Dryad repository.

## INTRODUCTION

Temperature impacts transmission of mosquito-borne diseases via effects on the physiology of mosquitoes and pathogens. Transmission requires that mosquitoes be abundant, bite a host and ingest an infectious bloodmeal, survive long enough for pathogen development and within-host migration (the extrinsic incubation period), and bite additional hosts—all processes that depend on temperature (Mordecai et al. 2013, 2017). Although both mechanistic (Mordecai et al. 2013, 2017, Liu-Helmersson et al. 2014, Wesolowski et al. 2015, Paull et al. 2017) and statistical models (Perkins et al. 2015, Siraj et al. 2015, Paull et al. 2017, Peña-García et al. 2017) support the impact of temperature on mosquito-borne disease, important knowledge gaps remain. First, how the impact of temperature on transmission differs across diseases, via what mechanisms, and the types of data needed to characterize these differences all remain uncertain. Second, the impacts of temperature on transmission can appear idiosyncratic—varying in both magnitude and direction—across locations and studies (Gatton et al. 2005, Stewart Ibarra and Lowe 2013, Peña-García et al. 2017). Although inferring causality from field observations and statistical approaches alone remains challenging, nonlinear thermal biology may mechanistically explain this variation. As the climate changes, filling these gaps becomes increasingly important for predicting geographic, seasonal, and interannual variation in transmission of mosquito-borne pathogens. Here, we address these gaps by building a model for temperature-dependent transmission of Ross River virus (RRV), the most important mosquito-borne disease in Australia (1,500–9,500 human cases per year) (Koolhof et al. 2017), and a potentially emerging pathogen worldwide (Flies et al. 2018).

RRV in Australia is an ideal case study for examining the influence of temperature. Transmission occurs across a wide latitudinal gradient, where climate varies substantially both geographically and seasonally. Moreover, compared to vector-borne diseases in lower-income settings, RRV case diagnosis and reporting are accurate and consistent, and variation in socioeconomic conditions (and therefore housing and vector control efforts) at regional and continental scales is relatively low. Previous work has shown that in some settings temperature predicts RRV cases (Gatton et al. 2005, Bi et al. 2009, Werner et al. 2012, Koolhof et al. 2017), while in others it does not (Hu et al. 2004, Gatton et al. 2005). Understanding RRV transmission ecology is critical because the virus is a likely candidate for emergence worldwide (Flies et al. 2018), and has caused explosive epidemics where it has emerged in the past (infecting over 500,000 people in a 1979-80 epidemic in Fiji) (Klapsing et al. 2005). RRV is a significant public health burden because infection causes joint pain that can become chronic and cause disability (Harley et al. 2001, Koolhof et al. 2017). A mechanistic model for temperature-dependent transmission could help explain these disparate results and predict potential expansion.

Mechanistic models synthesize how environmental factors like temperature influence host and parasite traits that drive transmission. Thermal responses of ectotherm traits are usually unimodal: they peak at intermediate temperatures and decline towards zero at lower and upper thermal limits, all of which vary across traits (Dell et al. 2011, Mordecai et al. 2013, 2017). Mechanistic models are particularly useful for synthesizing the effects of multiple, nonlinear thermal responses that shape transmission (Rogers and Randolph 2006, Mordecai et al. 2013). One commonly used measure of disease spread is *R_0_*, the basic reproductive number (secondary cases expected from a single case in a fully susceptible population). For mosquito-borne disease, *R_0_* is a nonlinear function of mosquito density, biting rate, vector competence (infectiousness given pathogen exposure), and adult survival; pathogen extrinsic incubation period; and human recovery rate (Dietz 1993). To understand how multiple traits that respond nonlinearly to temperature combine to affect transmission, we incorporate empirically-estimated trait thermal responses into an *R_0_* model. Synthesizing the full suite of nonlinear trait responses is critical because such models often make predictions that are drastically different, with transmission optima up to 7°C lower, than models that assume linear or monotonic thermal responses or omit temperature-dependent processes (Mordecai et al. 2013, 2017). Previous mechanistic models that incorporate multiple nonlinear trait thermal responses have predicted different optimal temperatures across pathogens and vector species: 25°C for *falciparum* malaria and West Nile virus (Mordecai et al. 2013, Paull et al. 2017), and 29°C and 26 °C for dengue, chikungunya, and Zika viruses in *Ae. aegypti* and *Ae. albopictus*, respectively (Liu-Helmersson et al. 2014, Wesolowski et al. 2015, Mordecai et al. 2017).

Here, we build the first mechanistic model for temperature-dependent transmission of RRV and ask whether temperature explains seasonal and geographic patterns of disease. We use data from laboratory experiments with two important and well-studied vector species (*Culex annulirostris* and *Aedes vigilax*) to parameterize the model with unimodal thermal responses. We then use sensitivity and uncertainty analyses to determine which traits drive the relationship between temperature and transmission potential and identify key data gaps. Finally, we illustrate how temperature currently shapes patterns of disease transmission across Australia. The model correctly predicts that RRV disease is year-round endemic in tropical, northern Australia with little seasonal variation due to temperature and seasonally epidemic in temperate, southern Australia. These results provides a mechanistic explanation for idiosyncrasies in RRV temperature responses observed in previous studies (Hu et al. 2004, Gatton et al. 2005, Bi et al. 2009, Werner et al. 2012, Koolhof et al. 2017). A population-weighted version of the model also accurately predicts the seasonality of human cases nationally. Thus, from laboratory data on mosquito and parasite thermal responses alone, this simple model mechanistically explains broad geographic and seasonal patterns of disease.

### Natural History of RRV

The natural history of RRV is complex: transmission occurs across a range of climates (tropical, subtropical, and temperate) and habitats (urban and rural, coastal and inland) and via many vector and vertebrate reservoir species (Claflin and Webb 2015). The virus has been isolated from over 40 mosquito species in nature, and 10 species transmit it in laboratory studies (Harley et al. 2001, Russell 2002). However, four species are responsible for most transmission to humans (*Culex annulirostris, Aedes [Ochlerotatus] vigilax, Ae. [O.] notoscriptus*, and *Ae. [O.] camptorhynchus*), with two additional species implicated in outbreaks (*Ae. [Stegomyia] polynesiensis* and *Ae. [O.] normanensis*).

The vectors differ in climate and habitat niches, leading to geographic variation in associations with outbreaks. We assembled and mapped records of RRV outbreaks in humans attributed to different vector species (Fig. 1, Table S1). *Ae. vigilax* and *Ae. notoscriptus* were more commonly implicated in transmission in tropical and subtropical zones, *Ae. camptorhynchus* in temperate zones, and *Cx. annulirostris* throughout all climatic zones. Freshwater-breeding *Cx. annulirostris* has been implicated in transmission across both inland and coastal areas, while saltmarsh mosquitoes *Ae. vigilax* and *Ae. camptorhynchus* have been implicated only in coastal areas (Russell 2002) and inland areas affected by salinization from agriculture (Biggs and Mottram 2008, Carver et al. 2009). Peri-domestic, container-breeding *Ae. notoscriptus* has been implicated in urban epidemics (Russell 2002). The vectors also differ in their seasonality: *Ae. camptorhynchus* populations peak earlier and in cooler temperatures than *Ae. vigilax*, leading to seasonal succession where they overlap (Russell 1998). This latitudinal and temporal variation suggests that vector species may have different thermal optima and/or niche breadths. If so, temperature may impact disease transmission differently for each species.

**FIGURE 1:**
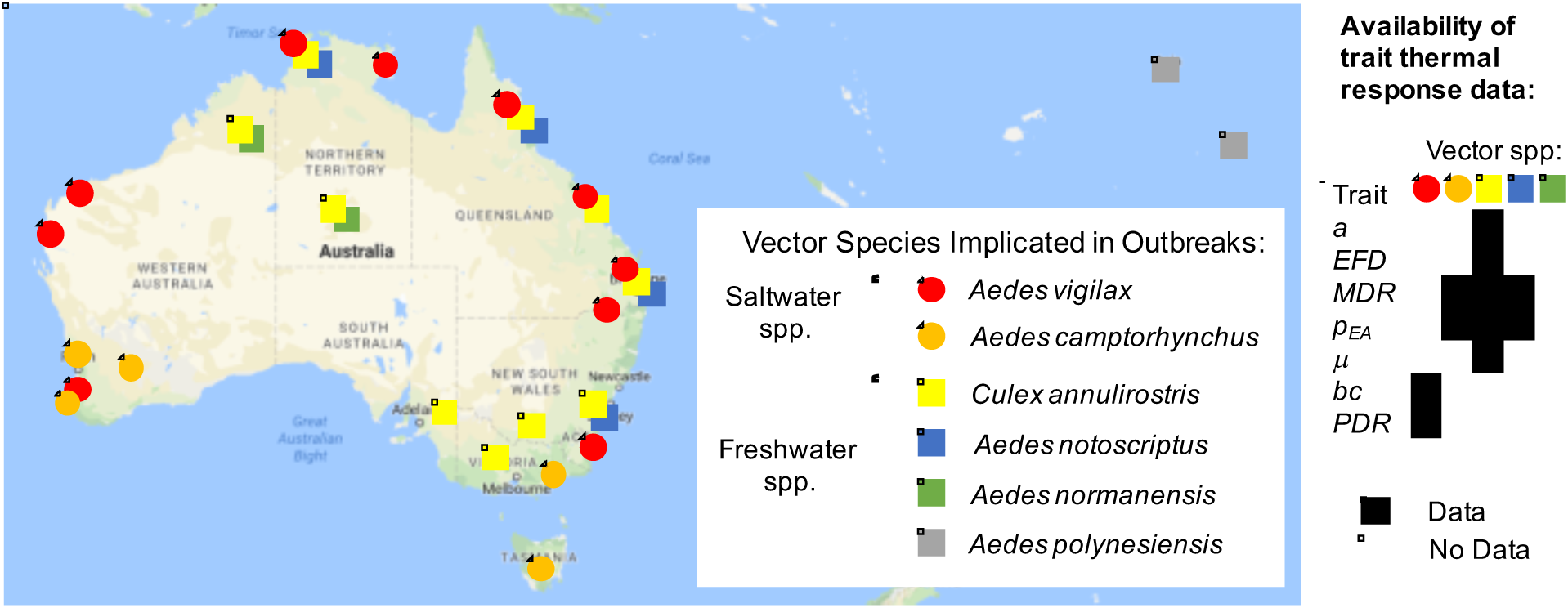
Vector species implicated in RRV disease outbreaks. Map of specific mosquito species identified as important vectors based on collected field specimens. Grid (right) shows data availability of trait thermal responses for the five Australian species. Data sources listed in Table S1. Trait parameters defined in Methods.

### General Modelling Approach

*R_0_* depends on a suite of vector, pathogen, and human traits, including vector density. Vector density depends on both temperature-dependent traits and aquatic breeding habitat (Barton et al. 2004, Kokkinn et al. 2009, Jacups et al. 2015). Therefore, we developed two *R_0_* models: the ‘Constant *M* Model’ (eq. 1) assumes mosquito density (*M*) does not depend on temperature (Dietz 1993); the ‘Temperature-Dependent *M* Model’ (eq. 2) assumes temperature drives mosquito density and includes vector life history trait thermal responses (Parham and Michael 2010, Mordecai et al. 2013, 2017). The relative influence of temperature versus habitat availability and other drivers varies across settings; these models capture two extremes. Because habitat availability is context-dependent and species-specific, and because we focus on temperature, we do not model vector habitat here. We initially compare results for both models, then focus on the temperature-dependent *M* model.

Since *R_0_* also depends on other factors, we scaled model output between zero and one (‘relative *R_0_*’). Relative *R_0_* describes thermal suitability for transmission, which combines with factors like breeding habitat availability, vector control, humidity, human and reservoir host density, host immune status, and mosquito exposure to determine disease incidence. In this approach, only the relative thermal response of each trait influences *R_0_*, desirable since traits can differ substantially due to other factors and in laboratory versus field settings (particularly mosquito survival: Clements and Paterson 1981). For relative *R_0_* we cannot use the typical threshold for sustained disease transmission (*R_0_*>1). However, relative *R_0_* preserves the temperature-dependence of *R_0_*, including key temperature values: where transmission is possible (*R_0_*>0; a conservative threshold where transmission is not excluded by temperature) and where *R_0_* is maximized.

## RESULTS

Vector and pathogen traits that drive transmission consistently responded to temperature (Fig. 2), though data were sparse (Table S2) (McDonald et al. 1980, Mottram et al. 1986, Russell 1986, Rae 1990, Kay and Jennings 2002). Although we exhaustively searched for experiments with trait measurements at three or more constant temperatures in the Australian vector species (*Cx. annulirostris, Ae. vigilax, Ae. camptorhynchus, Ae. notoscriptus*, and *Ae. normanensis*), no species had data for all necessary traits (Fig. 1). Thus, we combined mosquito life history traits from *Cx. annulirostris* and infection traits measured in *Ae. vigilax* to build composite *R_0_* models. We also fit traits for other mosquito and virus species: *MDR* and *pEA* from *Ae. camptorhynchus* and *Ae. notoscriptus*, and *PDR* and *bc* from Murray Valley encephalitis virus (another important pathogen transmitted by these mosquitoes in Australia) in *Cx. annulirostris* (see Appendix). We use sensitivity analyses to evaluate the potential impact of this vector mismatch.

**FIGURE 2:**
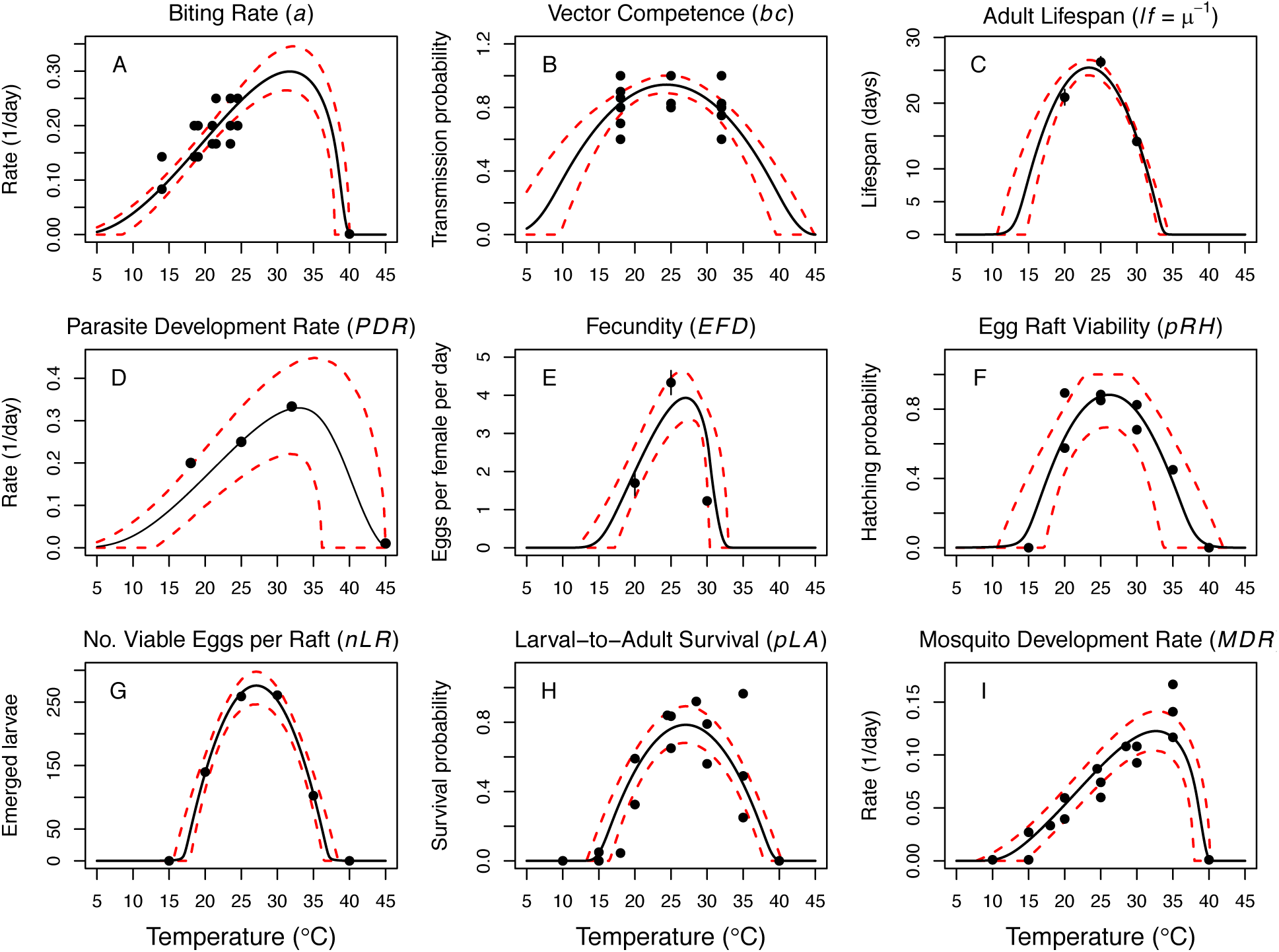
Thermal responses of *Cx. annulirostris* and RRV (in *Ae. vigilax*) traits that drive transmission. Functions were fit using Bayesian inference with priors fit using data from other mosquito species and viruses. Black solid lines are posterior distribution means; dashed red lines are 95% credible intervals. (E, C) Points are data means; error bars are standard error.

Thermal optima ranged from 23.4°C for adult lifespan (*lf*) to 33.0°C for parasite development rate (*PDR;* Fig. 2). The data supported unimodal thermal responses for most traits, though declines at high temperatures were not directly observed for biting rate (*a*) and parasite development rate. Instead, data from other mosquito species and ectotherm physiology theory imply these traits must decline at very high temperatures (~40°C).

The optimal temperature for transmission (relative *R_0_*) peaked near 26.4°C regardless of whether or not mosquito density [*M*] depended on temperature, because optimal transmission aligned with optimal mosquito density (Fig. 3; temperature-dependent *M* model: 26.4°C, constant *M* model: 26.6°C, *M* 26.2°C). By contrast, the range of temperatures suitable for transmission is much narrower when mosquito density depends on temperature because *M*(*T*) constrains transmission at the thermal limits (Fig. 3; temperature-dependent *M* model: 17.0-31.5°C, constant *M* model: 12.9–33.7°C). The thermal constraints that mosquito density imposes on transmission are important because, although demographic traits are well-known to vary with temperature in the laboratory, many temperature-dependent transmission models do not assume that temperature influences mosquito density (Martens et al. 1997, Craig et al. 1999, Caminade et al. 2017, Paull et al. 2017, Hamlet et al. 2018, but see Parham and Michael 2010, Mordecai et al. 2013, 2017, Johnson et al. 2015). The moderate optimal temperature for RRV (26-27°C) fits within the range of thermal optima found for other diseases: malaria transmission by *Anopheles* spp. at 25°C, and dengue and other viruses by *Ae. aegypti* and *Ae. albopictus* at 29°C and 26°C, respectively (Fig. 4) (Mordecai et al. 2013, 2017).

**FIGURE 3:**
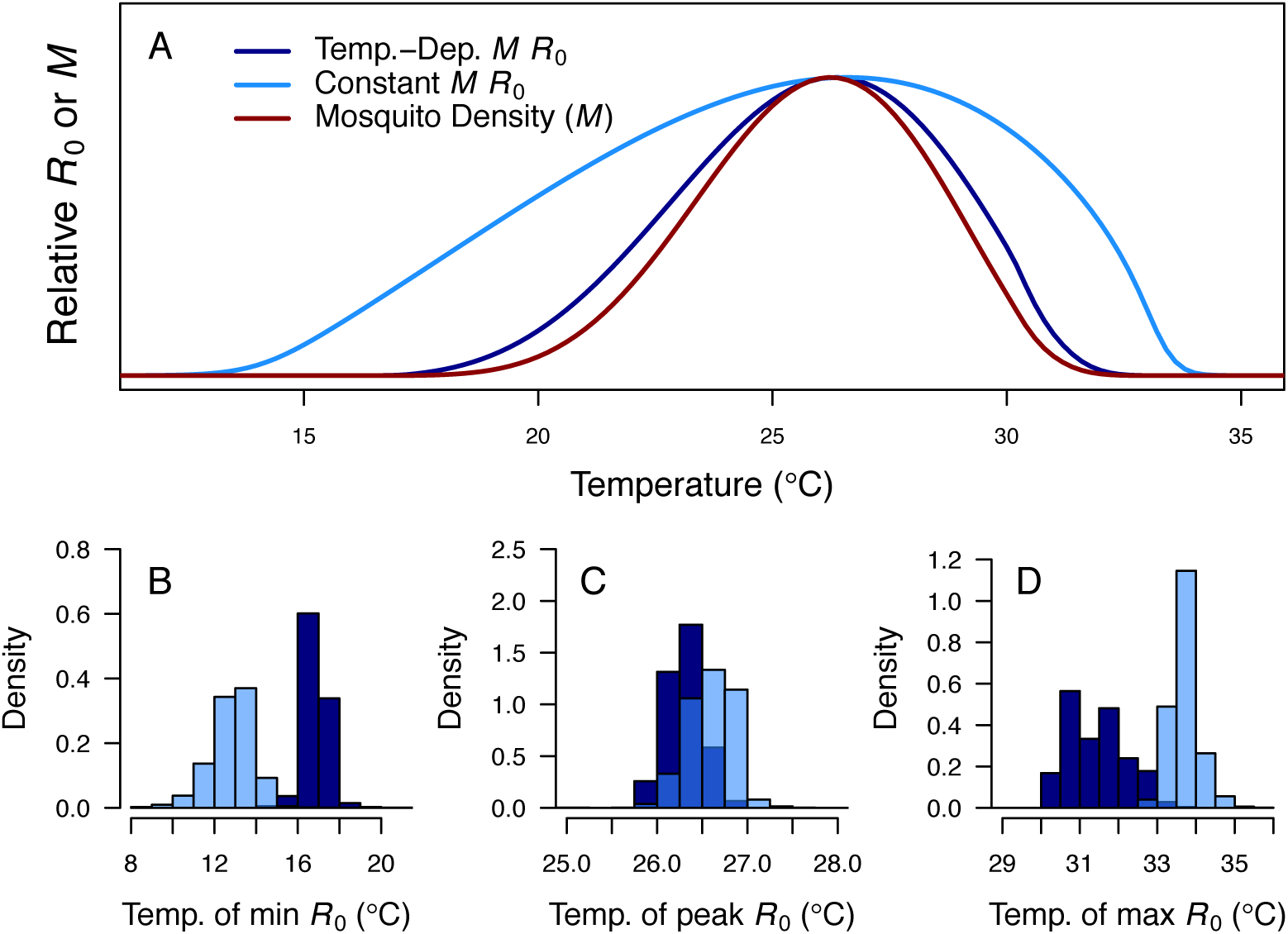
Thermal response of relative *R_0_*. (A) Posterior means across temperature for constant *M* model (eq. 1, light blue) and temperature-dependent *M* model (eq. 2, dark blue). Predicted mosquito density (*M*) shown for comparison (red). The y-axis shows relative *R_0_* (or *M*) rather than absolute values, which would require additional information. Histograms of (B) the critical thermal minimum, (C) thermal optimum, and (D) critical thermal maximum temperatures for both models (same colors as in A).

**FIGURE 4:**
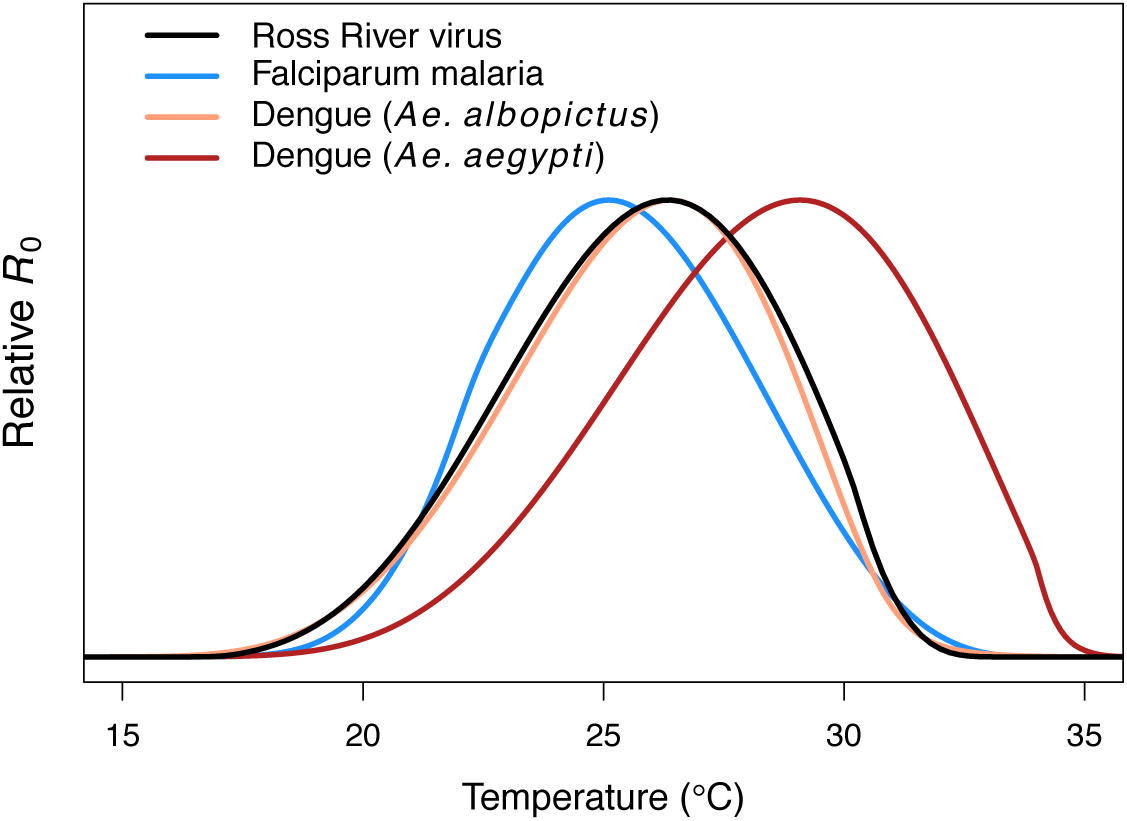
Comparing the RRV *R_0_* model to other diseases: malaria (blue, optimum = 25°C), Ross River virus (black, optimum = 26.4°C), dengue virus in *Ae. albopictus* (orange, optimum = 26.4°C), and dengue virus in *Ae. agypti* (red, optimum = 29.1°C).

At the upper thermal limit fecundity (*EFD*) and adult lifespan (*lf*) constrain *R_0_*, while at the lower thermal limit fecundity, larval survival (*pLA*), egg survival (raft viability [*pRH*] and survival within rafts [*nLR*]), and adult lifespan constrain *R_0_* (Fig. S6). All of these traits (except adult lifespan) only occur in, and adult lifespan is quantitatively more important in, the temperature-dependent *M* model. Correspondingly, uncertainty in these traits generated the most uncertainty in *R_0_* at the respective thermal limits (Fig. S6C). The optimal temperature for *R_0_* was most sensitive to the thermal response of adult lifespan. Near the optimum, most uncertainty in *R_0_* was due to uncertainty in the thermal responses of adult lifespan, egg raft viability, and fecundity. Substituting larval traits from alternative vectors or infection traits for Murray Valley Encephalitis virus did not substantially alter the *R_0_* thermal response, since *Cx. annulirostris* life history traits strongly constrained transmission (Fig. S5).

Temperature suitability for RRV transmission varies seasonally across Australia, based on the temperature-dependent *M* model (eq. 2) using monthly mean temperatures from WorldClim. In subtropical and temperate locations (Brisbane and further south), low temperatures force *R_0_* to zero for part of the year (Figs. 5A, 6). Monthly mean temperatures in these areas fall along the increasing portion of the *R_0_* curve for the entire year, so thermal suitability for transmission increases with temperature. By contrast, in tropical, northern Australia (Darwin and Cairns), the temperature remains suitable throughout the year (Figs. 5, 6). Darwin is the only major city where mean temperatures exceed the thermal optimum, and thereby depress transmission. Because most Australians live in southern, temperate areas, country-scale transmission is seasonal. As hypothesized *a priori*, human cases peak two months after population-weighted *R_0_*(*T*) (Fig. 7).

**FIGURE 5:**
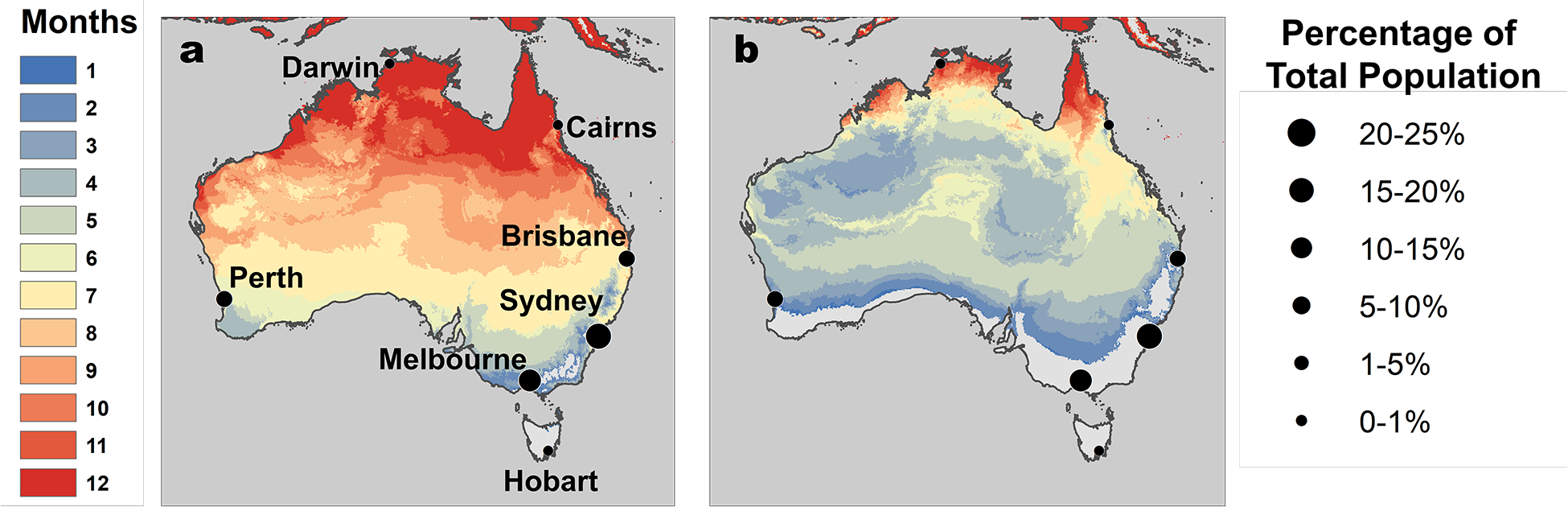
RRV transmission potential from monthly mean temperatures. Color indicates number of months where (A) relative *R_0_*>0 and (B) relative *R_0_*>0.5. Predictions are based on the posterior distribution medians. Points indicate selected cities (Fig. 5), scaled by the percent of total Australian population residing in each city.

**FIGURE 6:**
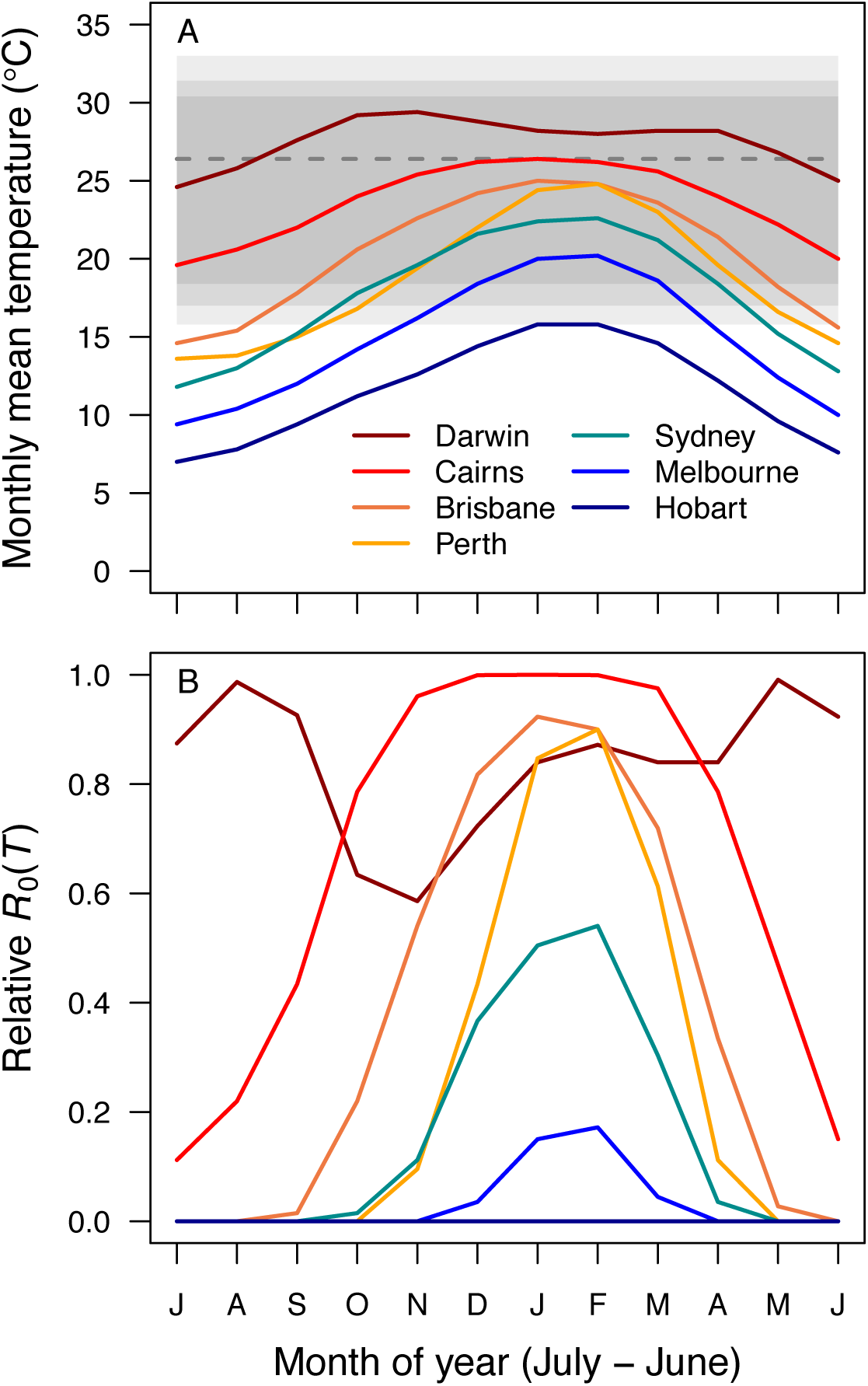
Average seasonality of temperature and relative *R_0_* in Australian cities. The selected cities span a latitudinal and temperature gradient (Darwin = dark red, Cairns = red, Brisbane = dark orange, Perth = light orange, Sydney = aqua, Melbourne = blue, Hobart = dark blue). The x-axis begins in July and ends in June (during winter). (A) Mean monthly temperatures. Shaded areas show temperature thresholds where *R_0_*>0 for: outer 95% CI (light grey), median (medium grey), and inner 95% CI (dark grey). Dashed line shows median *R_0_* optimal temperature. (B) Temperature-dependent *R_0_*.

**Figure 7:**
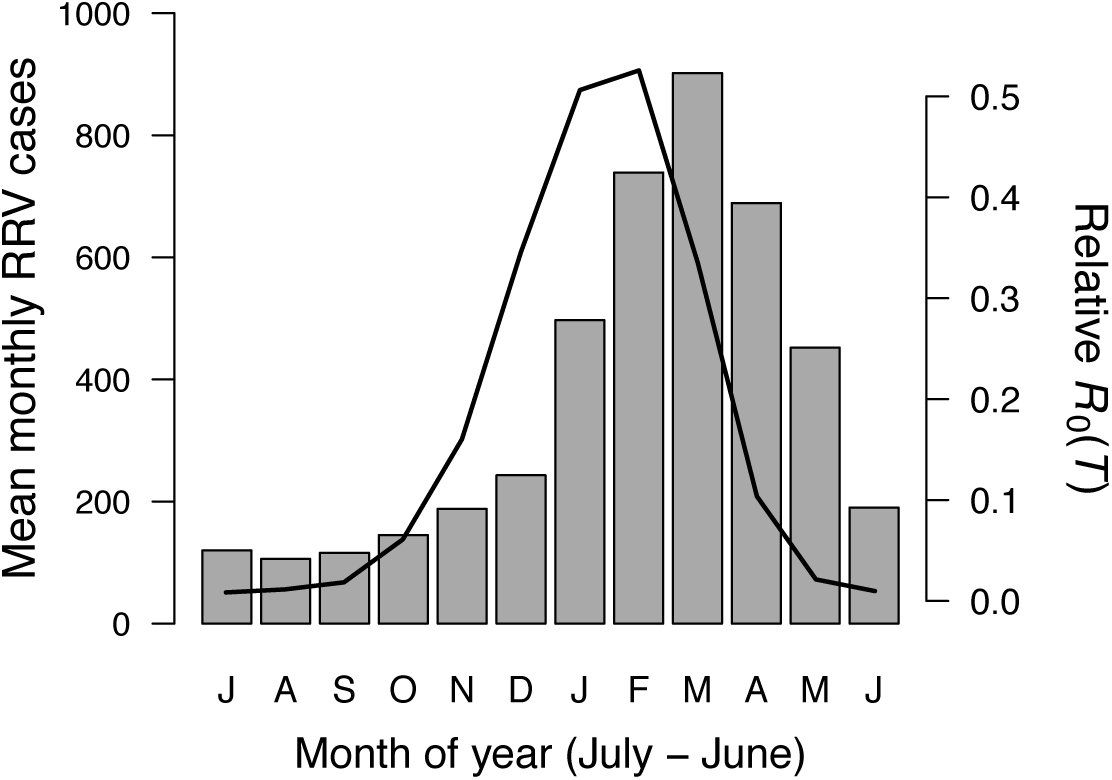
Seasonality of relative *R_0_* and RRV infections. Human cases aggregated nationwide from 1992-2013 (bars). Temperature-dependent *R_0_* weighted by population (line), calculated from Australia’s 15 largest cities (76.6% of total population). The x-axis begins in July and ends in June (during winter). Cases peak two months after *R_0_*, the *a priori* expected lag between temperature and reported cases.

## DISCUSSION

In a warming world, it is critical to understand effects of temperature on transmission of mosquito-borne disease, particularly as new mosquito-borne pathogens emerge and spread worldwide. Identifying transmission optima and limits by characterizing nonlinear thermal responses, rather than simply assuming that transmission increases with temperature, can more accurately predict geographic, seasonal, and interannual variation in disease. Thermal responses vary substantially among diseases and vector species (Mordecai et al. 2013, 2017), yet we lack mechanistic models based on empirical, unimodal thermal responses for many diseases and vectors. Here, we parameterized a temperature-dependent model for transmission of RRV (Fig. 2) with data from two important vector species (*Cx. annulirostris* and *Ae. vigilax*; Fig. 1). The optimal temperature for transmission is moderate (26-27°C; Fig. 3), and largely determined by the thermal response of adult mosquito lifespan (Fig. S6). Both low and high temperatures limit transmission due to low mosquito fecundity and survival at all life stages (not just adults)—thermal responses that are often ignored in transmission models (Fig. S6). Temperature explains the geography of year-round endemic versus seasonally epidemic disease (Figs. 5, 6) and accurately predicts the seasonality of human cases at the national scale (Fig. 7). Thus, the model for RRV transmission provides a mechanistic link between geographic and seasonal variation in temperature and broad-scale patterns of disease.

While the thermal response of RRV transmission generally matched those of other mosquito-borne pathogens, there were some key differences. The moderate optimal temperature for RRV (26-27°C) fit within the range of thermal optima found for other diseases: malaria transmission by *Anopheles* spp. at 25°C, and dengue and other viruses by *Ae. aegypti* and *Ae. albopictus* at 29°C and 26°C, respectively (Fig. 4) (Mordecai et al. 2013, 2017). For all of these diseases, the specific temperature for optimal transmission was largely determined by the thermal response of adult lifespan (Mordecai et al. 2013, 2017, Johnson et al. 2015). However, the traits that set the thermal limits for RRV transmission differed from other systems. The lower thermal limit for RRV was constrained by fecundity and survival at all stages while the upper thermal limit was constrained by fecundity and adult lifespan. By contrast, thermal limits for malaria transmission were set by parasite development rate at cool temperatures and egg-to-adulthood survival at high temperatures (Mordecai et al. 2013). As with previous models, the upper and lower thermal limits of RRV transmission are more uncertain than the optimum (Fig. 3) (Johnson et al. 2015, Mordecai et al. 2017), because trait responses are harder to measure near their thermal limits where survival is low and development is slow or incomplete. Overall, our results support a general pattern of intermediate thermal optima for transmission where the well-resolved optimal temperature is driven by adult mosquito lifespan, but upper and lower thermal limits are more uncertain and may be determined by unique traits for different vectors and pathogens.

The trait thermal response data were limited in two keys ways. First, two traits (fecundity and adult lifespan) had data from only three temperatures. We used priors derived from data from other mosquito species to minimize over-fitting and better represent the true fit and uncertainty (Fig. 2, versus uniform priors in Fig. S1). However, data from more temperatures would increase our confidence in the fitted thermal responses. Second, no vector species had data for all traits (Fig. 1), so we combined mosquito traits from *Cx. annulirostris* and pathogen infection traits in *Ae. vigilax* to build composite *R_0_* models. Geographic and seasonal variation in vector populations suggests that *Ae. camptorhynchus* and *Ae. vigilax* have different thermal niches (cooler and warmer, respectively) and *Cx. annulirostris* has a broader thermal niche (Fig. 1) (Russell 1998). We need temperature-dependent trait data for more species to test the hypothesis that these niche differences reflect the species’ thermal responses. If true, the current model, parameterized primarily with *Cx. annulirostris* trait responses, may not accurately predict transmission by *Ae. camptorhynchus* and *Ae. vigilax*. Hypothesized species differences in thermal niche could explain why RRV persists over a wide climatic and latitudinal gradient. Thus, thermal response experiments with other RRV vectors are a critical area for future research.

The temperature-dependent *R_0_* model provides a mechanistic explanation for independently-observed patterns of RRV transmission across Australia. As predicted (Figs. 5, 6), RRV is endemic in tropical Australia, with little seasonal variation in transmission potential due to temperature, and seasonally epidemic in subtropical and temperate Australia (Weinstein 1997). The model also accurately predicts disease seasonality at the national scale (Fig. 7), reproducing the *a priori* predicted lag (8-10 weeks) for temperature to affect reported human cases (Hu et al. 2006, Jacups et al. 2008, Stewart Ibarra et al. 2013, Mordecai et al. 2017). Further, RRV transmission by *Cx. annulirostris* in inland areas often moves south as temperatures increase from spring into summer (Russell 1998), matching the model prediction (Fig. 6). Although temperature is often invoked as a potential driver for such patterns, it is difficult to establish causality from statistical inference alone, particularly if temperature and disease both exhibit strong seasonality and could both be responding to another latent driver. Thus, the mechanistic model is a critical piece of evidence linking temperature to patterns of disease.

In addition to explaining broad-scale patterns, the unimodal thermal model explains previously contradictory local-scale results. Specifically, statistical evidence for temperature impacts on local time series of cases is mixed. RRV incidence is often—but not always—positively associated with warmer temperatures (Tong and Hu 2001, Tong et al. 2002, 2004, Hu et al. 2004, 2010, Jacups et al. 2008, Williams et al. 2009, Werner et al. 2012, Koolhof et al. 2017). However, variation in temperature impacts across space and time is expected from an intermediate thermal optimum, especially when observed mean temperatures are near or cross the optimum. The strongest statistical signal of temperature on disease is expected in temperate regions where mean temperature varies along the rapidly rising portion of the *R_0_* curve (~20-25°C). If mean temperatures vary both above and below the optimum (as in Darwin), important effects of temperature may be masked in time series models that fit linear responses.

Additionally, if temperatures are always relatively suitable (as in tropical climates) or unsuitable (as in very cool temperate climates), variation in disease may be due primarily to other factors. A nonlinear mechanistic model is critical for estimating temperature impacts on transmission because the effect of increasing temperature by a few degrees can have a positive, negligible, or negative impact on *R_0_* along different parts of the thermal response curve. Although field-based evidence for unimodal thermal responses in vector-borne disease is rare (but see Mordecai et al. 2013, Perkins et al. 2015, Peña-García et al. 2017), there is some evidence for high temperatures constraining RRV transmission and vector populations: outbreaks were less likely with more days above 35°C in part of Queensland (Gatton et al. 2005) and populations of *Cx. annulirostris* peaked at 25°C and declined above 32°C in Victoria (Dhileepan 1996). Future statistical analyses of RRV cases may benefit from using a nonlinear function for temperature-dependent *R_0_* as a predictor instead of raw temperature (Fig. 5B versus 5A).

Breeding habitat availability also drives mosquito abundance and mosquito-borne disease. Local rainfall and river flow have been linked to the abundance of RRV vector species (Tall et al. 2014) and RRV disease cases (Tong and Hu 2001, 2002a, Hu et al. 2004, Kelly-Hope et al. 2004, Tong et al. 2004, Gatton et al. 2005, Jacups et al. 2008, Bi et al. 2009, Williams et al. 2009, Werner et al. 2012), as have unusually high tides in coastal areas with saltmarsh mosquitoes (Tong and Hu 2002b, Tong et al. 2004, Jacups et al. 2008). Overlaying models of species-specific breeding habitat with temperature-dependent models will better resolve the geographic and seasonal distribution of RRV transmission. *R_0_* peaked at similar temperatures whether or not we assumed mosquito abundance was temperature-dependent (eq. 1 versus eq. 2); however, the range of suitable temperatures was much wider for the model that assumed a temperature-independent mosquito population (Fig. 3). Since breeding habitat can only impact vector populations when temperatures do not exclude them, it is critical to consider thermal constraints on mosquito abundance, even when breeding habitat is considered a stronger driver. Nonetheless, many mechanistic, temperature-dependent models of vector-borne disease transmission do not include thermal effects on vector density (Martens et al. 1997, Craig et al. 1999, Caminade et al. 2017, Paull et al. 2017, Hamlet et al. 2018). Our results demonstrate that the decision to exclude these relationships can have a critical impact on model results, especially near thermal limits.

Several important gaps remain in our understanding of RRV thermal ecology, in addition to the need for trait thermal response data for more vector species. First, the *R_0_* model needs to be more rigorously validated using time series of cases to determine the importance of temperature at finer spatiotemporal scales. These analyses should incorporate daily and seasonal temperature variation (Paaijmans et al. 2010) and integrate species-specific drivers of breeding habitat availability, like rainfall and tidal patterns. Second, translating environmental suitability for transmission into human cases also depends on disease dynamics in reservoir host populations and their impact on immunity. For instance, in Western Australia heavy summer rains can fail to initiate epidemics when low rainfall in the preceding winter depresses recruitment of susceptible juvenile kangaroos (Mackenzie et al. 2000). By contrast, large outbreaks occur in southeastern Australia when high rainfall follows a dry year, presumably from higher transmission within relatively unexposed reservoir populations (Woodruff et al. 2002). Building vector species-specific *R_0_* models and integrating thermal ecology with other drivers are important next steps for forecasting variation in RRV transmission.

Nonlinear thermal responses are particularly important for predicting how transmission will change under future climate regimes. Climate warming will likely increase the geographic and seasonal range of transmission potential in temperate, southern Australia where most Australians live. However, climate change will likely decrease transmission potential in tropical areas like Darwin, where moderate warming (~3°C) would push temperatures above the upper thermal limit for transmission for most of the year (Fig. 5). However, the extent of climate-driven declines in transmission will depend on how much *Cx. annulirostris* and *Ae. vigilax* can adapt to extend their upper thermal limits and whether warmer-adapted vector species (e.g., *Ae. aegypti* and potentially *Ae. polynesiensis*) can invade and sustain RRV transmission cycles. Thus, we can predict the response of RRV transmission by current vector species to climate change based on these trait thermal responses. However, future disease dynamics will also depend on vector adaptation, potential vector species invasions, and climate change impacts on sea level and precipitation that drive vector habitat availability.

## METHODS

### Temperature-Dependent R_0_ Models

We used two *R_0_* models: ‘Constant *M* Model’ (eq. 1) assumes mosquito density (*M*) does not depend on temperature (equation from Dietz 1993); ‘Temperature-Dependent *M* Model’ (eq. 2) assumes temperature drives mosquito density and includes vector life history trait thermal responses (Parham and Michael 2010, Mordecai et al. 2013, 2017).

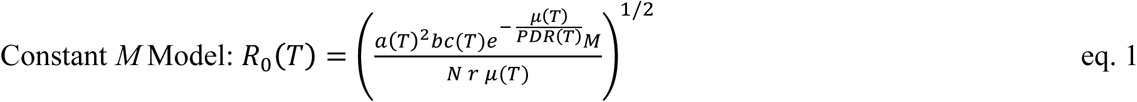

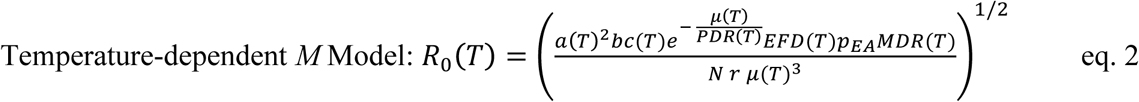

In both equations, (*T*) indicates a parameter depends on temperature, *a* is mosquito biting rate, *bc* is vector competence (proportion of mosquitoes becoming infectious post-exposure), *μ* is adult mosquito mortality rate (adult lifespan, *lf* = 1/*μ*), *PDR* is parasite development rate (*PDR* = 1/*EIP*, the extrinsic incubation period), *N* is human density, and *r* is the recovery rate at which humans become immune (all rates are measured in days^-1^). The latter two terms do not depend on temperature. In the temperature-dependent *M* model, mosquito density (*M*) depends on fecundity (*EFD*, eggs per female per day), proportion surviving from egg-to-adulthood (*p_EA_*), and mosquito development rate (*MDR*), divided by the square of adult mortality rate (*μ*) (Parham and Michael 2010). We calculated *p_EA_* as the product of the proportion of egg rafts that hatch (*pRH*), the number of larvae per raft (*nLR*, scaled by the maximum at any temperature to calculate proportional egg survival within-rafts), and the proportion of larvae surviving to adulthood (*p_LA_*).

We fit thermal responses of traits in the *R_0_* models to previously published data (Table S2) using Bayesian inference with the ‘r2jags’ package (Plummer 2003, Su and Yajima 2009) in R (R Core Team 2017). Traits with asymmetrical thermal responses were fit as Brière functions: *qT*(*T–T_0_*)(*T_max_–T*)^1/2^ (Brière et al. 1999). Traits with symmetrical thermal responses were fit as quadratic functions: -*q*(*T–T_0_*)(*T–T_max_*). In both functions of temperature (*T*), *T_0_* and *T_max_* are the critical thermal minimum and maximum, respectively, and *q* is a rate parameter. For priors we used gamma distributions with hyperparameters derived from thermal responses fit to data from other mosquito species (Table S4), allowing us to more accurately represent the fit and uncertainty. Our data did not include declining trait values at high temperatures for biting rate (*a*) and parasite development rate (*PDR*). Nonetheless, data from other mosquito species (Mordecai et al. 2013, 2017) and principles of thermal biology (Dell et al. 2011) imply these traits must decline at high temperatures. Thus, for those traits we included an artificial data point where the trait value approached zero at a very high temperature (40°C), allowing us to fit the Brière function. We used strongly informative priors to limit the effect of these traits on the upper thermal limit of *R_0_* (by constraining them to decline near 40°C). For comparison, we also fit all thermal responses with uniform priors (Fig. S1); these results illustrate how the priors impacted the results.

### Sensitivity and uncertainty analyses

We conducted sensitivity and uncertainty analyses of the temperature-dependent *M* model (eq. 2) to understand how trait thermal responses shape the thermal response of *R_0_*. We examined the sensitivity of *R_0_* two ways. First, we evaluated the impact of each trait by setting it constant while allowing all other traits to vary with temperature. Second, we calculated the partial derivative of *R_0_* with respect to each trait across temperature (*∂R_0_/∂X · ∂X/∂T* for trait *X* and temperature *T*; Appendix). To understand what data would most improve the model, we also calculated the proportion of total uncertainty in *R_0_* due to each trait across temperature. First, we propagated posterior samples from all trait thermal response distributions through to *R_0_*(*T*) and calculated the width of the 95% highest posterior density interval (HPD interval; a type of credible interval) of this distribution at each temperature: the ‘full *R_0_*(*T*) uncertainty’. Next, we sampled each trait from its posterior distribution while setting all other trait thermal responses to their posterior medians, and calculated the posterior distribution of *R_0_*(*T*) and the width of its 95% HPD interval across temperature: the ‘single-trait *R_0_*(*T*) uncertainty’. Finally, we divided each single-trait *R_0_*(*T*) uncertainty by the full *R_0_*(*T*) uncertainty.

### Field Observations: Seasonality of Temperature-Dependent R_0_ Across Australia

We took monthly mean temperatures from WorldClim for seven cities spanning a latitudinal and temperature gradient (from tropical North to temperate South: Darwin, Cairns, Brisbane, Perth, Sydney, Melbourne, and Hobart) and calculated the posterior median *R_0_*(*T*) for each month at each location. We also compared the seasonality of a population-weighted *R_0_*(*T*) and nationally aggregated RRV cases. We used 2016 estimates for the fifteen most populous urban areas, which together contain 76.6% of Australia’s population (Australian Bureau of Statistics 2017). We calculated *R_0_*(*T*) for each location (as above) and estimated a population-weighted average. We compared this country-scale estimate of *R_0_*(*T*) with data on mean monthly human cases of RRV nationwide from 1992-2013 obtained from the National Notifiable Diseases Surveillance System.

We expected a time lag between temperature and reported human cases as mosquito populations increase, bite humans and reservoir hosts, acquire RRV, become infectious, and bite subsequent hosts; after an incubation period hosts (potentially) become symptomatic, seek treatment, and report cases. Empirical work on dengue vectors in Ecuador identified a six-week time lag between temperature and mosquito oviposition (Stewart Ibarra et al. 2013). Subsequent mosquito development and incubation periods in mosquitoes and humans likely add another 2-4 week lag before cases appear, resulting in an 8-10 week lag between temperature and observed cases (Hu et al. 2006, Jacups et al. 2008, Mordecai et al. 2017). With monthly case data, we hypothesize a two-month time lag between *R_0_*(*T*) and RRV disease cases.

### Mapping Temperature-Dependent R_0_ Across Australia

To illustrate temperature suitability for RRV transmission across Australia, we mapped the number of months for which relative *R_0_*(*T*)>0 and >0.5 for the posterior median, 2.5%, and 97.5% credibility bounds (Fig. S7) for the temperature-dependent *M* model (eq. 2). We calculated *R_0_*(*T*) at 0.2°C increments and projected it onto the landscape for monthly mean temperatures from WorldClim data at a 5-minute resolution (approximately 10km^2^ at the equator). Climate data layers were extracted for the geographic area, defined using the Global Administrative Boundaries Databases (GADM 2012). We performed map calculations and manipulations in R with packages ‘raster’ (Hijmans 2016), ‘maptools’ (Bivand and Lewin-Koh 2017), and ‘Rgdal’ (Bivand et al. 2017), and rendered GeoTiffs in ArcGIS version 10.3.1.

## ACKNOWLEDGEMENTS

We thank Leah Johnson and Matt Thomas for comments and Cameron Webb for posting the RRV case data (https://cameronwebb.wordpress.com/2014/04/09/why-is-mosquito-borne-disease-risk-greater-in-autumn/).

## Appendix S1 Additional Notes, Tables, Figures, Methods, and References

### Notes

Notes on (1) mosquito nomenclature, (2) the use of medians versus means in the trait and *R_0_* thermal responses, and (3) data digitization.

#### Mosquito nomenclature

In 2000, there was a proposed shift in mosquito taxonomy: several subgenera within the genus *Aedes* were elevated to genus status (Wilkerson *et al*. 2015). This move affected the two saltmarsh mosquitoes included in this study, *Aedes vigilax* and *Aedes camptorhynchus*, which for a time were called *Ochlerotataus vigilax* and *Ochlerotatus camptorhynchus* by some researchers. More recently, there has been a consensus to move back to the previous naming system, so we use *Aedes* here, although many of the papers we cite use *Ocherlotatus* instead.

#### The use of means versus medians in thermal responses

Bayesian inference produces output in the form of posterior distributions rather than a single estimated value. Because these distributions can be non-normal and asymmetric, best practice is to report medians rather than means, since medians are less sensitive to outlying values in extended tails. In this study, we report median parameter values and use median predicted values for downstream analyses (e.g., sensitivity and uncertainty analyses, predictions using weather data). However, we plot mean values in the figures because they show a smoother and more visually intuitive representation of where trait and *R_0_* thermal responses go to zero at the upper thermal limits. The means and medians are not substantially different, except at these thermal limits (Fig S8).

#### Data digitization

We digitized previously published data using a free web-based tool called Webplot digitizer available at: https://automeris.io/WebPlotDigitizer/.

**TABLE S1:**
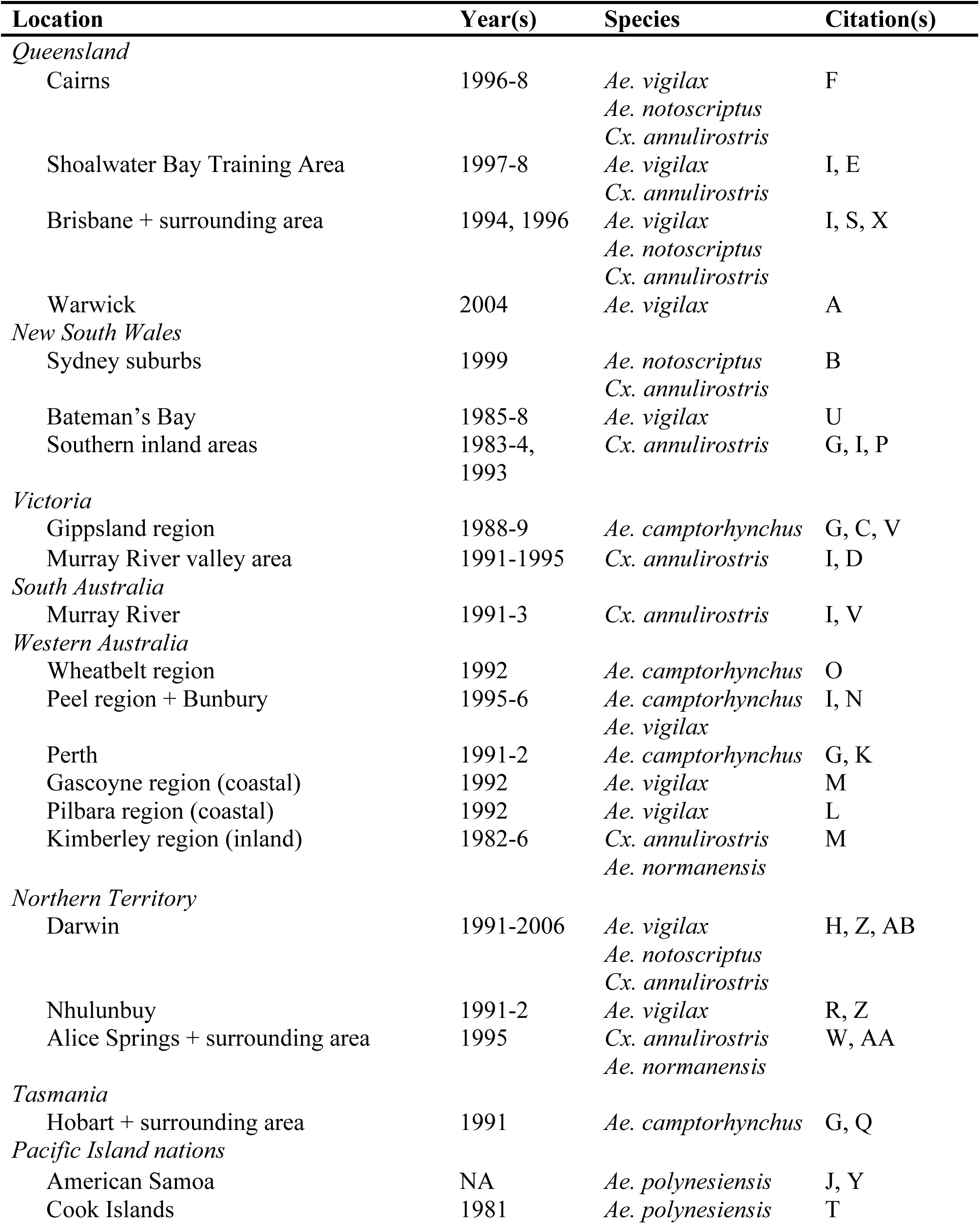
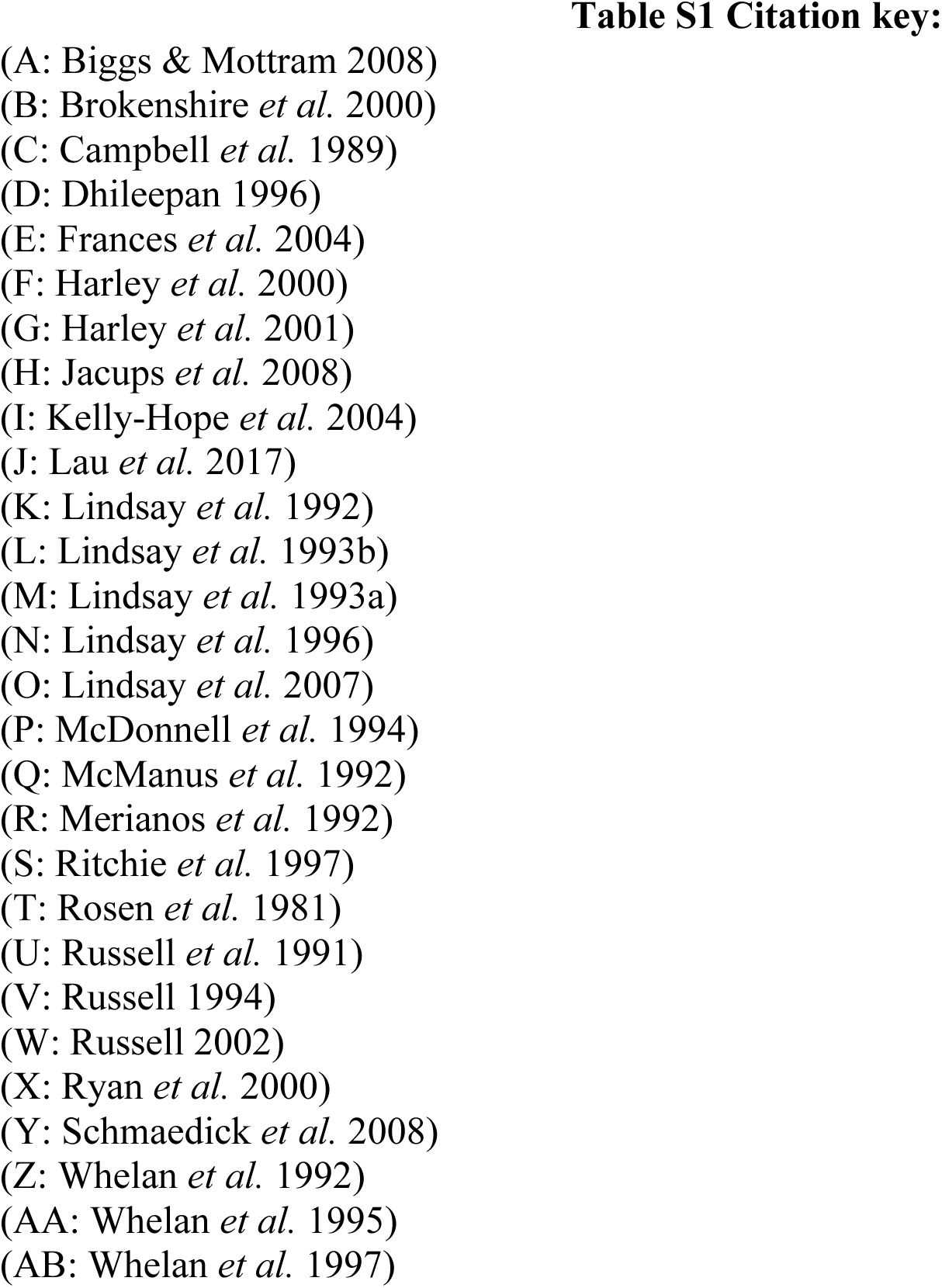
Mosquito species implicated in RRV outbreaks. (Fig. 1). Location and year of RRV disease outbreaks and mosquito species identified as likely vectors based on the collection of field specimens. Only the six most important mosquito species are included.

**TABLE S2:** Trait thermal response functions and data sources for Ross River virus *R_0_* models (eqs. 1 and 2). ‘Par.’ = model parameter. Results are given for fits from data-informed priors. Asymmetrical responses fit with Brière function (**B**): B(*T*) = *qT*(*T–T_0_*)(*T_max_–T*)^1/2^; symmetrical responses fit with quadratic function (**Q**): Q(*T*) = -*q*(*T–T_0_*)(*T–T_max_*). Function coefficients (and 95% credible intervals) fit via Bayesian inference.

| Par. | Definition | Species (Sources) | Fit | Function Coefficients (95% CIs) & Optimal Temperature |
| --- | --- | --- | --- | --- |
| $a$ | Biting rate = 1 / gonotrophic cycle duration (day)<sup>-1</sup> | *Cx. annulirostris* (Russell 1986) | **B** | $T_0 = 4.0$ (0.7 – 8.5)<br>$T_{max} = 39.1$ (37.9 – 40.0)<br>$q = 1.26 \cdot 10^{-4}$ (1.02 – $1.70 \cdot 10^{-4}$)<br>optimum = 31.8°C |
| $bc$ | Vector competence (transmission probability) | *Ae. vigilax* (Kay & Jennings 2002) | **Q** | $T_0 = 6.0$ (1.6 – 9.6)<br>$T_{max} = 42.7$ (39.6 – 44.9)<br>$q = 2.88 \cdot 10^{-3}$ (2.02 – $4.27 \cdot 10^{-3}$)<br>optimum = 24.4°C |
| $lf = 1/\mu$ | Adult lifespan (days) | *Cx. annulirostris* (McDonald *et al.* 1980) | **Q** | $T_0 = 13.0$ (10.6 – 14.6)<br>$T_{max} = 33.6$ (33.0 – 34.5)<br>$q = 0.241$ (0.177 – 0.304)<br>optimum = 23.4°C |
| $PDR$ | Parasite development rate (day)<sup>-1</sup> | *Ae. vigilax* (Kay & Jennings 2002) | **B** | $T_0 = 6.3$ (1.3 – 13.0)<br>$T_{max} = 41.1$ (36.1 – 45.0)<br>$q = 1.38 \cdot 10^{-4}$ (0.855 – $2.14 \cdot 10^{-4}$)<br>optimum = 33.0°C |
| EFD | Fecundity (eggs per female per day) | *Cx. annulirostris* (McDonald *et al.* 1980) | **Q** | $T_0 = 14.7$ (11.9 – 17.2)<br>$T_{max} = 31.4$ (30.3 – 32.9)<br>$q = 5.84 \cdot 10^{-3}$ (3.90 – $8.71 \cdot 10^{-3}$)<br>optimum = 27.0°C |
| pRH | Raft viability (probability of raft hatching) | *Cx. annulirostris* (McDonald *et al.* 1980; Mottram *et al.* 1986) | **Q** | $T_0 = 14.3$ (10.6 – 17.2)<br>$T_{max} = 38.1$ (33.7 – 42.0)<br>$q = 6.78 \cdot 10^{-3}$ (3.36 – $12.6 \cdot 10^{-3}$)<br>optimum = 26.2°C |
| nLR | Within-raft egg survival (number of larvae per raft) | *Cx. annulirostris* (Mottram *et al.* 1986) | **Q** | $T_0 = 17.0$ (15.6 – 17.9)<br>$T_{max} = 37.2$ (36.4 – 38.4)<br>$q = 2.73 \cdot 10^{-3}$ (1.99 – $3.29 \cdot 10^{-3}$)<br>optimum = 27.0°C |
| $pLA$ | Larval-to-adult survival (probability) | *Cx. annulirostris* (McDonald *et al.* 1980; Mottram *et al.* 1986; Rae 1990) | **Q** | $T_0 = 14.9$ (13.3 – 16.5)<br>$T_{max} = 39.1$ (37.9 – 40.4)<br>$q = 5.39 \cdot 10^{-3}$ (4.31 – $6.65 \cdot 10^{-3}$)<br>optimum = 27°C |
| $MDR$ | Mosquito development rate (day)<sup>-1</sup> | *Cx. annulirostris* (McDonald *et al.* 1980; Mottram *et al.* 1986; Rae 1990) | **B** | $T_0 = 11.1$ (7.6 – 14.4)<br>$T_{max} = 39.2$ (37.9 – 40.1)<br>$q = 6.86 \cdot 10^{-5}$ (5.47 – $8.47 \cdot 10^{-5}$)<br>optimum = 32.6°C |

**TABLE S3:** Trait thermal response functions and data sources for Murray Valley Encephalitis virus and additional vector species (*Ae. notoscriptus* and *Ae. camptorhynchus*). ‘Par.’ = model parameter. Results are given for fits from data-informed priors. Asymmetrical ½ responses fit with Brière function (**B**): B(*T*) = *qT*(*T–T_0_*)(*T_max_–T*)^1/2^; symmetrical responses fit with quadratic function (**Q**): Q(*T*) = -*q*(*T–T_0_*)(*T–T_max_*). Function coefficients (and 95% credible intervals) fit via Bayesian inference.

| Par. | Definition | Species (Sources) | Fit | Function Coefficients (95% CIs) & Optimal Temperature |
| --- | --- | --- | --- | --- |
| *bc* | Vector competence (transmission probability) | MVEV in *Cx. Annulirostris* (Kay *et al.* 1989) | NA | Did not fit because no temperature signal |
| *PDR* | Parasite development rate (day)<sup>-1</sup> | MVEV in *Cx. Annulirostris* (Kay *et al.* 1989) | **B** | $T_0 = 12.8$ (6.6 – 19.0)<br>$T_{max} = 41.9$ (38.6 – 45.0)<br>$q = 1.04 \cdot 10^{-4}$ (0.641 – $1.55 \cdot 10^{-4}$)<br>optimum = 34.8°C |
| *pLA* | Larval-to-adult survival (probability) | *Ae. camptorhynchus* (Barton & Aberton 2005) | **Q** | $T_0 = 5.5$ (1.9 – 10.1)<br>$T_{max} = 38.6$ (35.7 – 42.3)<br>$q = 3.02 \cdot 10^{-3}$ (2.00 – $4.83 \cdot 10^{-3}$)<br>optimum = 22.2°C |
| *pLA* | Larval-to-adult survival (probability) | *Ae. notoscriptus* (Williams & Rau 2011) | **Q** | $T_0 = 9.1$ (7.1 – 11.0)<br>$T_{max} = 36.2$ (34.7 – 37.9)<br>$q = 5.72 \cdot 10^{-3}$ (4.37 – $7.32 \cdot 10^{-3}$)<br>optimum = 22.8°C |
| *MDR* | Mosquito development rate (day)<sup>-1</sup> | *Ae. camptorhynchus* (Barton & Aberton 2005) | **B** | $T_0 = 9.5$ (1.8 – 19.1)<br>$T_{max} = 38.8$ (37.2 – 40.3)<br>$q = 4.57 \cdot 10^{-5}$ (2.50 – $8.29 \cdot 10^{-5}$)<br>optimum = 32.2°C |
| *MDR* | Mosquito development rate (day)<sup>-1</sup> | *Ae. notoscriptus* (Williams & Rau 2011) | **B** | $T_0 = 9.6$ (6.4 – 12.9)<br>$T_{max} = 38.7$ (37.1 – 40.0)<br>$q = 6.86 \cdot 10^{-5}$ (5.43 – $8.56 \cdot 10^{-5}$)<br>optimum = 32.0°C |

**TABLE S4:**
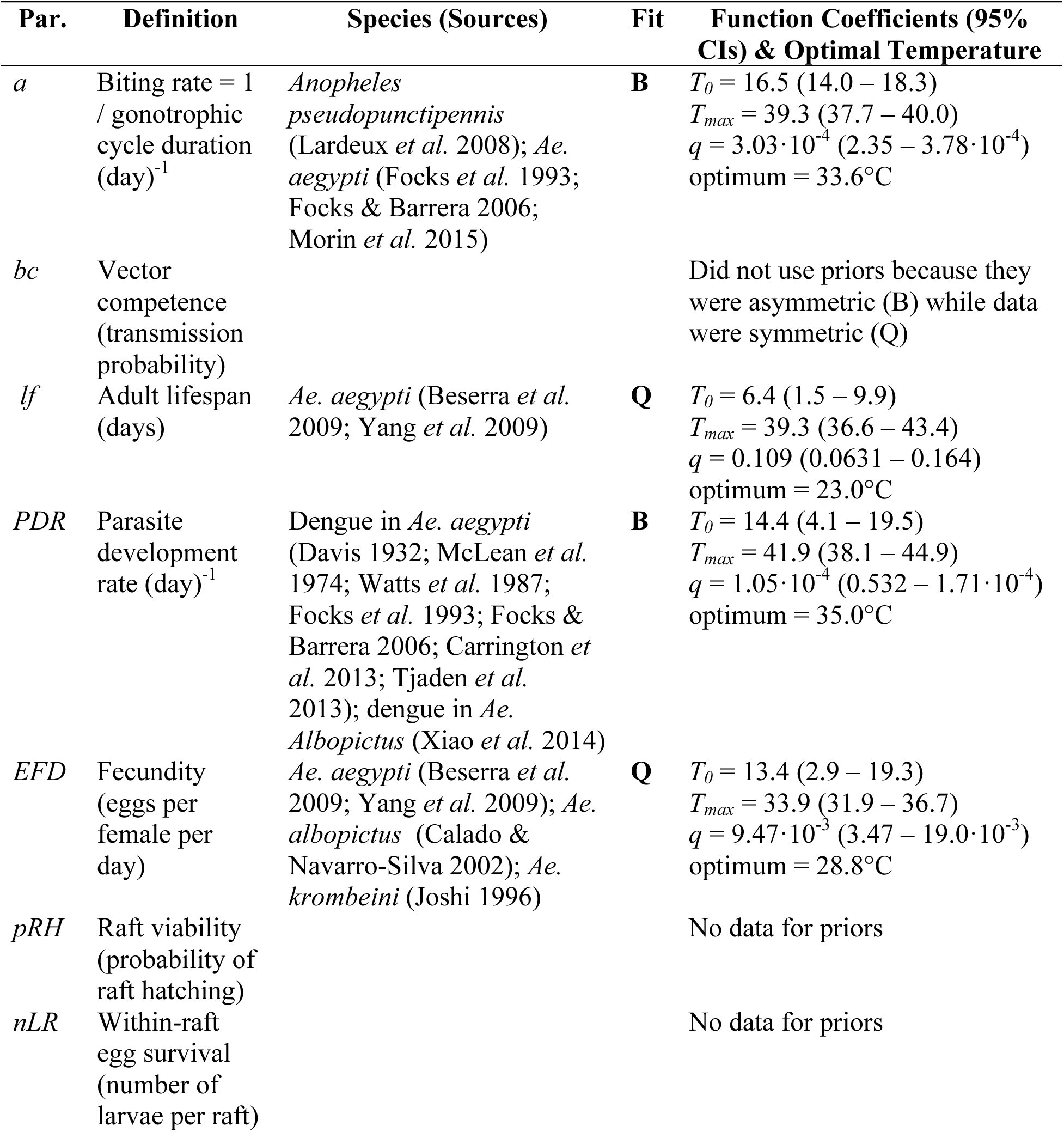
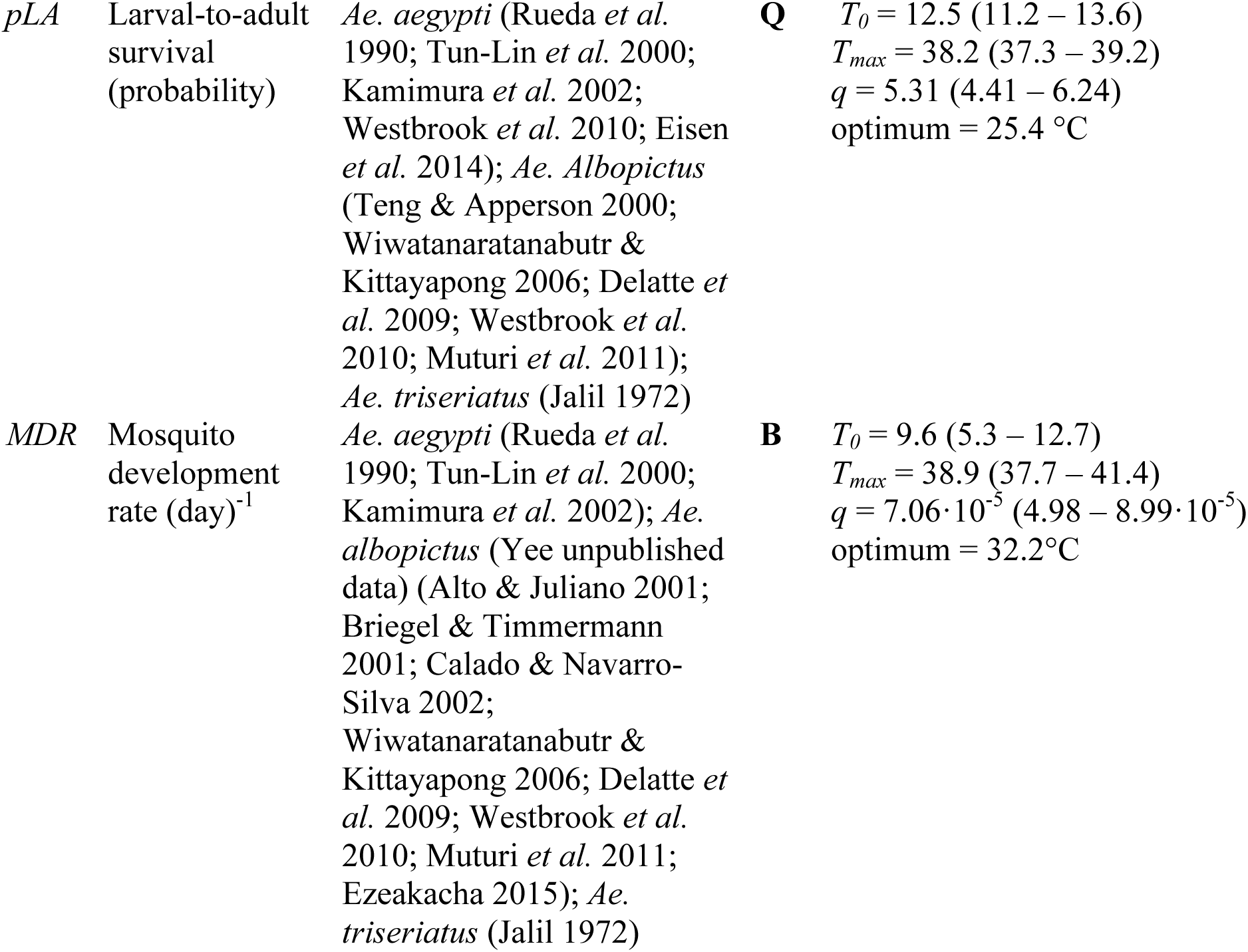
Trait thermal response functions and data sources used to parameterize priors for data-informed trait thermal responses. ‘Par.’ = model parameter. Fits were made with uniform priors. Asymmetrical responses fit with Brière function (**B**): B(*T*) = *qT*(*T–T_0_*)(*T_max_–T*)^1/2^; symmetrical responses fit with quadratic function (**Q**): Q(*T*) = -*q*(*T–T_0_*)(*T–T_max_*). Function coefficients (and 95% credible intervals) fit via Bayesian inference.

| Par. | Definition | Species (Sources) | Fit | Function Coefficients (95% CIs) & Optimal Temperature |
| --- | --- | --- | --- | --- |
| *a* | Biting rate = 1 / gonotrophic cycle duration (day)<sup>-1</sup> | *Anopheles pseudopunctipennis* (Lardeux *et al.* 2008); *Ae. aegypti* (Focks *et al.* 1993; Focks & Barrera 2006; Morin *et al.* 2015) | **B** | $T_0 = 16.5$ (14.0 – 18.3)<br>$T_{max} = 39.3$ (37.7 – 40.0)<br>$q = 3.03 \cdot 10^{-4}$ (2.35 – $3.78 \cdot 10^{-4}$)<br>optimum = 33.6°C |
| *bc* | Vector competence (transmission probability) |  |  | Did not use priors because they were asymmetric (B) while data were symmetric (Q) |
| *lf* | Adult lifespan (days) | *Ae. aegypti* (Beserra *et al.* 2009; Yang *et al.* 2009) | **Q** | $T_0 = 6.4$ (1.5 – 9.9)<br>$T_{max} = 39.3$ (36.6 – 43.4)<br>$q = 0.109$ (0.0631 – 0.164)<br>optimum = 23.0°C |
| *PDR* | Parasite development rate (day)<sup>-1</sup> | Dengue in *Ae. aegypti* (Davis 1932; McLean *et al.* 1974; Watts *et al.* 1987; Focks *et al.* 1993; Focks & Barrera 2006; Carrington *et al.* 2013; Tjaden *et al.* 2013); dengue in *Ae. Albopictus* (Xiao *et al.* 2014) | **B** | $T_0 = 14.4$ (4.1 – 19.5)<br>$T_{max} = 41.9$ (38.1 – 44.9)<br>$q = 1.05 \cdot 10^{-4}$ (0.532 – $1.71 \cdot 10^{-4}$)<br>optimum = 35.0°C |
| *EFD* | Fecundity (eggs per female per day) | *Ae. aegypti* (Beserra *et al.* 2009; Yang *et al.* 2009); *Ae. albopictus* (Calado & Navarro-Silva 2002); *Ae. krombeini* (Joshi 1996) | **Q** | $T_0 = 13.4$ (2.9 – 19.3)<br>$T_{max} = 33.9$ (31.9 – 36.7)<br>$q = 9.47 \cdot 10^{-3}$ (3.47 – $19.0 \cdot 10^{-3}$)<br>optimum = 28.8°C |
| *pRH* | Raft viability (probability of raft hatching) |  |  | No data for priors |
| *nLR* | Within-raft egg survival (number of larvae per raft) |  |  | No data for priors |
| *pLA* | Larval-to-adult survival (probability) | *Ae. aegypti* (Rueda *et al.* 1990; Tun-Lin *et al.* 2000; Kamimura *et al.* 2002; Westbrook *et al.* 2010; Eisen *et al.* 2014); *Ae. Albopictus* (Teng & Apperson 2000; Wiwatanaratanabutr & Kittayapong 2006; Delatte *et al.* 2009; Westbrook *et al.* 2010; Muturi *et al.* 2011); *Ae. triseriatus* (Jalil 1972) | **Q** | $T_0 = 12.5$ (11.2 – 13.6)<br>$T_{max} = 38.2$ (37.3 – 39.2)<br>$q = 5.31$ (4.41 – 6.24)<br>optimum = 25.4 °C |
| *MDR* | Mosquito development rate (day)<sup>-1</sup> | *Ae. aegypti* (Rueda *et al.* 1990; Tun-Lin *et al.* 2000; Kamimura *et al.* 2002); *Ae. albopictus* (Yee unpublished data) (Alto & Juliano 2001; Briegel & Timmermann 2001; Calado & Navarro-Silva 2002; Wiwatanaratanabutr & Kittayapong 2006; Delatte *et al.* 2009; Westbrook *et al.* 2010; Muturi *et al.* 2011; Ezeakacha 2015); *Ae. triseriatus* (Jalil 1972) | **B** | $T_0 = 9.6$ (5.3 – 12.7)<br>$T_{max} = 38.9$ (37.7 – 41.4)<br>$q = 7.06 \cdot 10^{-5}$ (4.98 – 8.99 · 10<sup>-5</sup>)<br>optimum = 32.2°C |

**FIGURE S1:**
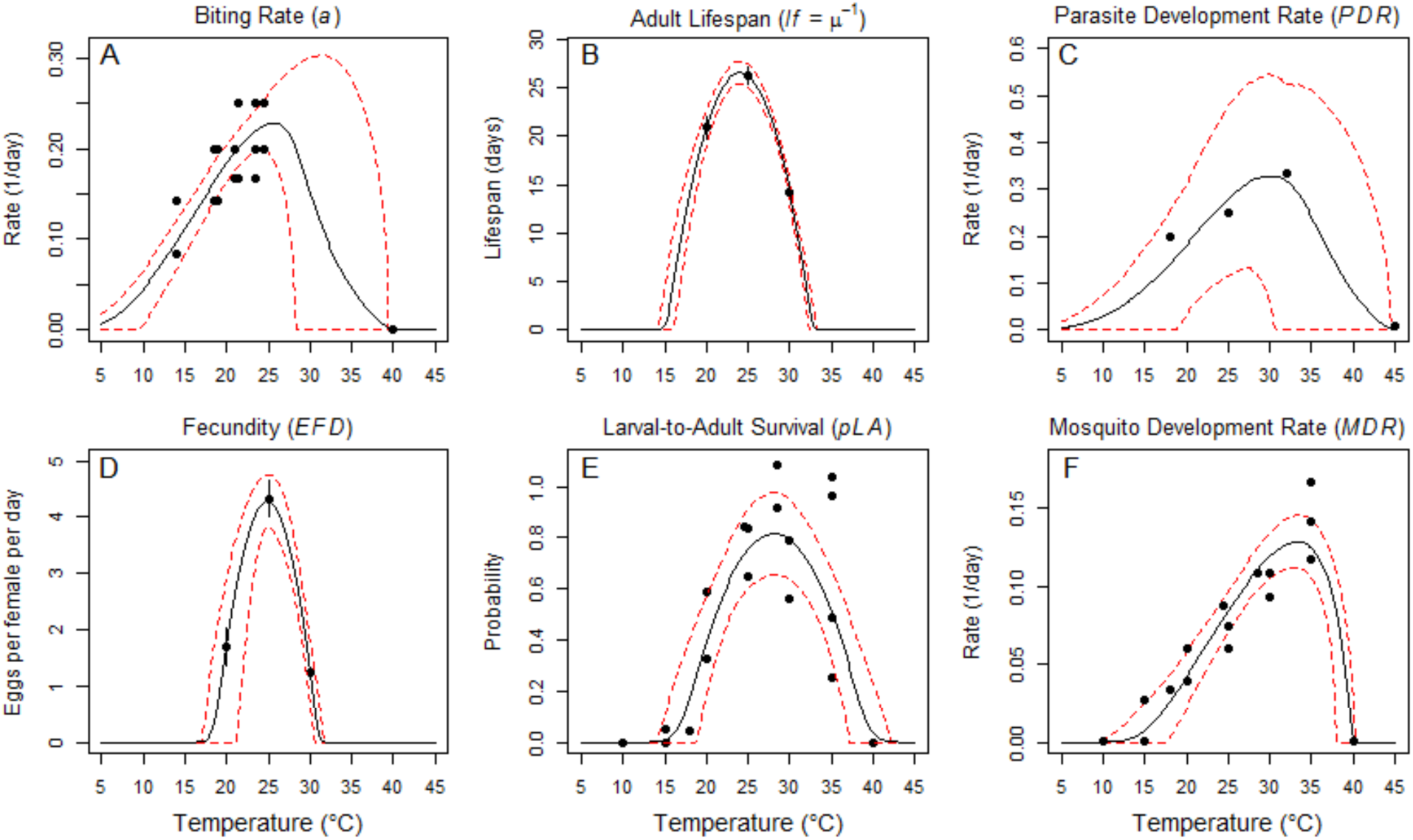
Thermal responses of *Cx. annulirostris* and Ross River virus (in *Ae. vigilax*) traits fit with uniform priors. For lifespan (*lf*; B) and fecundity (*EFD;* D) points show data means and error bars indicate standard error (for display only; models were fit to raw data). Black solid lines are function means; dashed red lines are function 95% credible intervals. The thermal responses for vector competence (*bc*) and egg survival traits (*pRH* and *nLR*) fit with uniform priors are included in Fig. 2 (main text) because we did not use data-informed priors for those traits.

**FIGURE S2:**
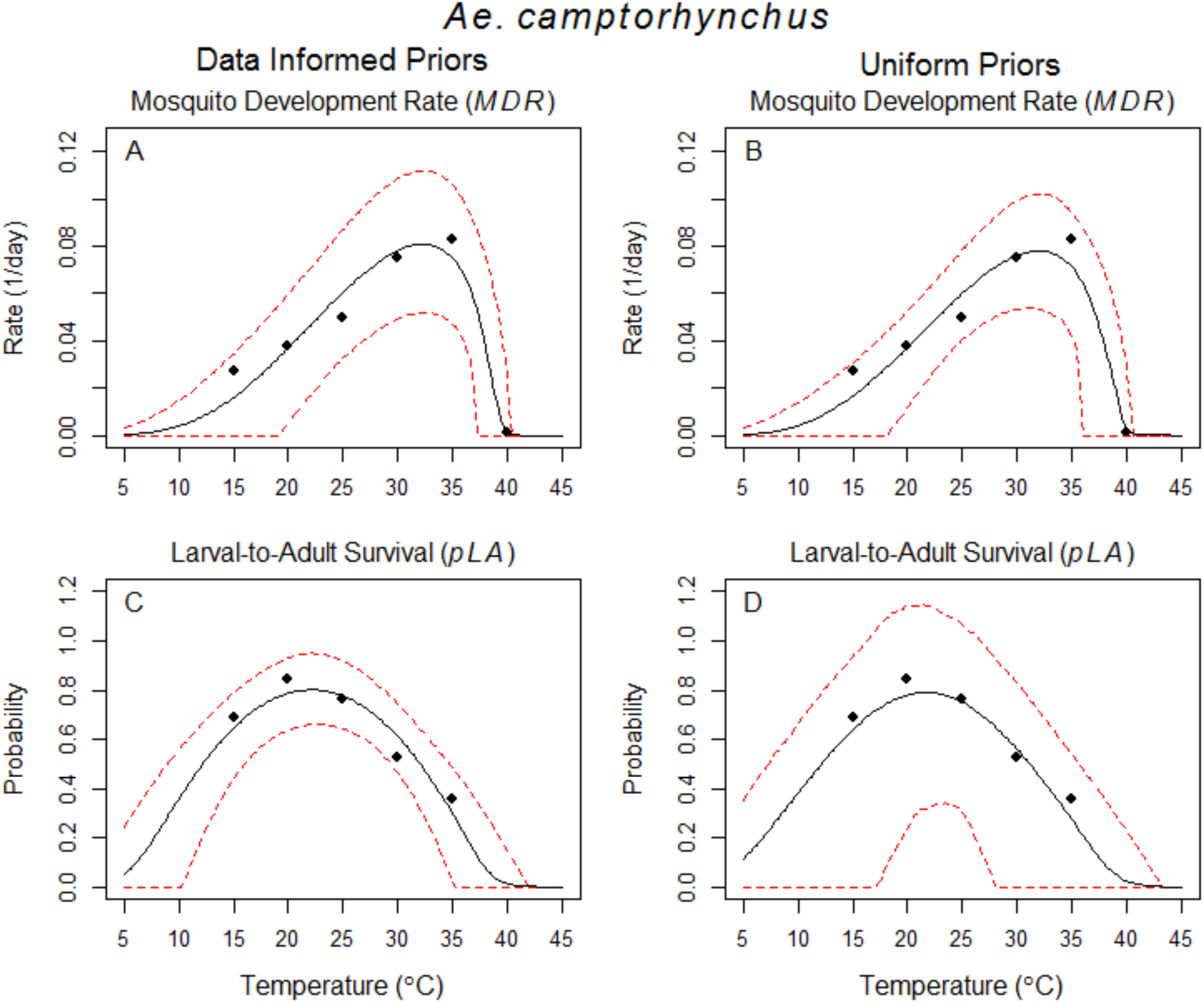
Trait thermal responses for *Ae. camptorhynchus*. (Top row) Mosquito development rate (*MDR*) and (bottom row) larval survival (*pLA*) fit with (left column) data-informed priors and (right column) uniform priors. Black solid lines are function means; dashed red lines are function 95% credible intervals.

**FIGURE S3:**
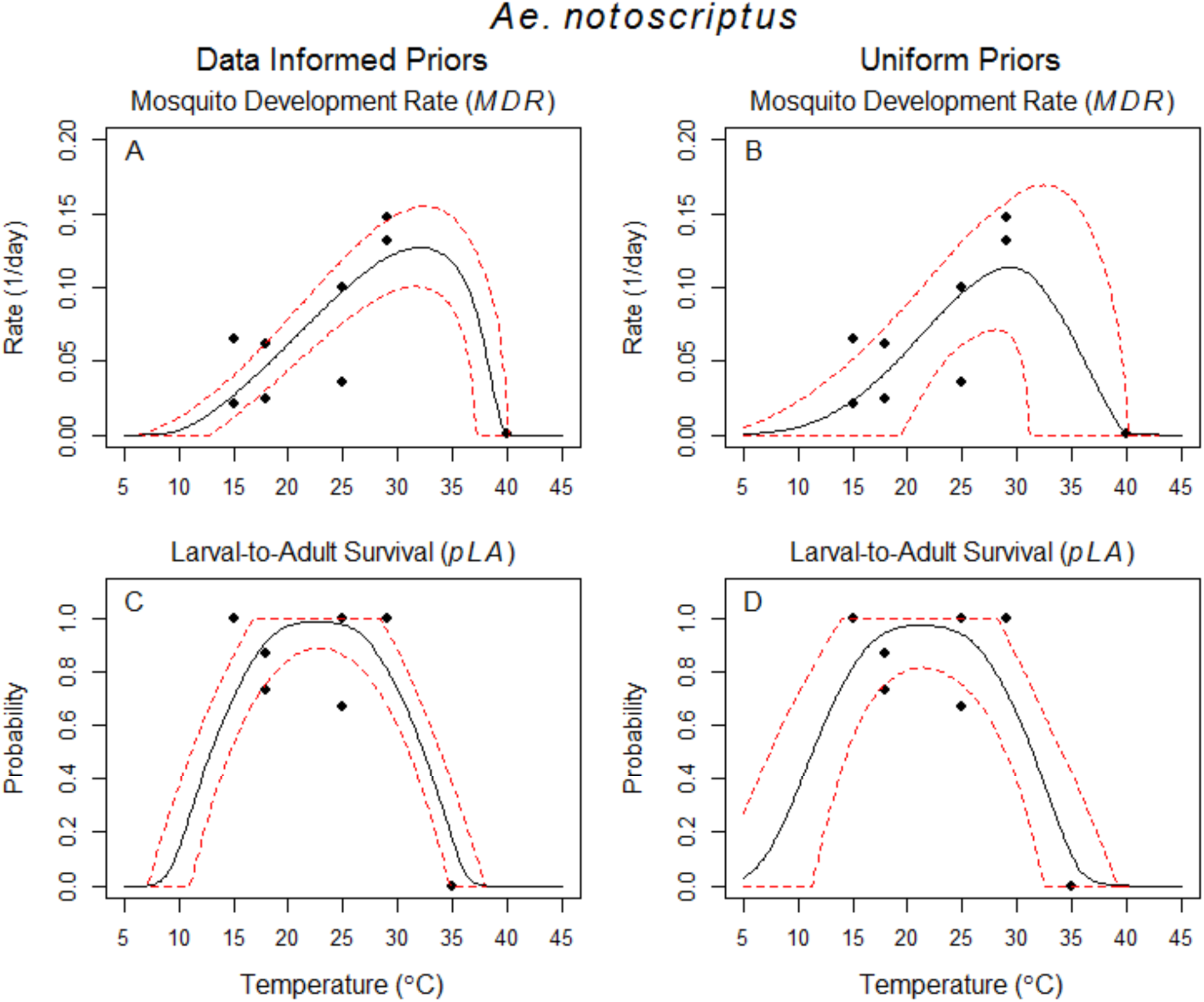
Trait thermal responses for *Ae. notoscriptus*. (Top row) Mosquito development rate (*MDR*) and (bottom row) larval survival (*pLA*) fit with (left column) data-informed priors and (right column) uniform priors. Black solid lines are function means; dashed red lines are function 95% credible intervals.

**FIGURE S4:**
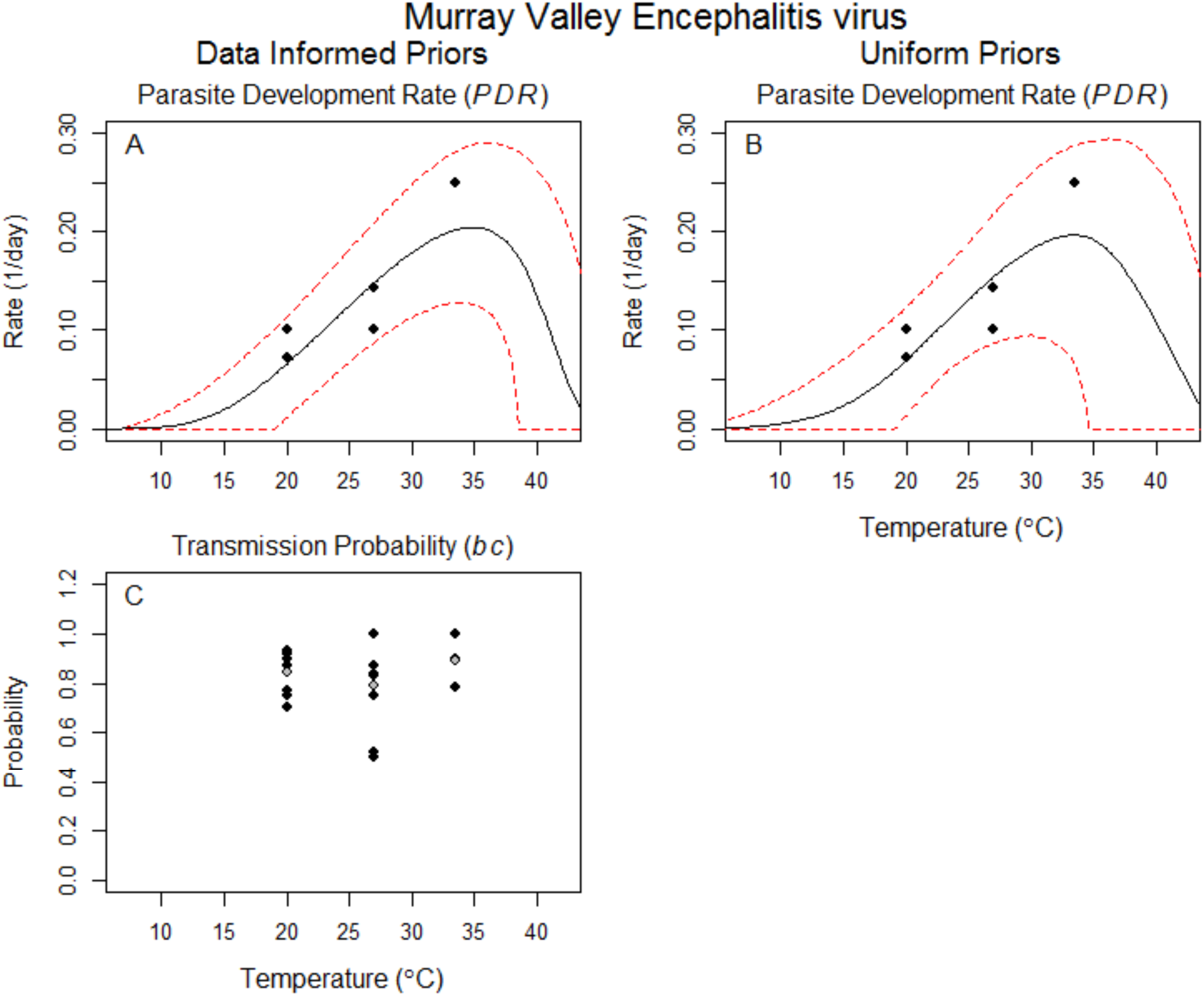
Trait thermal responses for Murray Valley Encephalitis virus in *Cx. annulirostris*. (Top row) Pathogen development rate (*PDR*) fit with (A) data-informed priors and (B) uniform priors. Black solid lines are function means; dashed red lines are function 95% credible intervals. (C) Data for vector competence (*bc*). Gray points are temperature treatment means. Because there was no temperature signal, we did not fit a thermal response.

**FIGURE S5:**
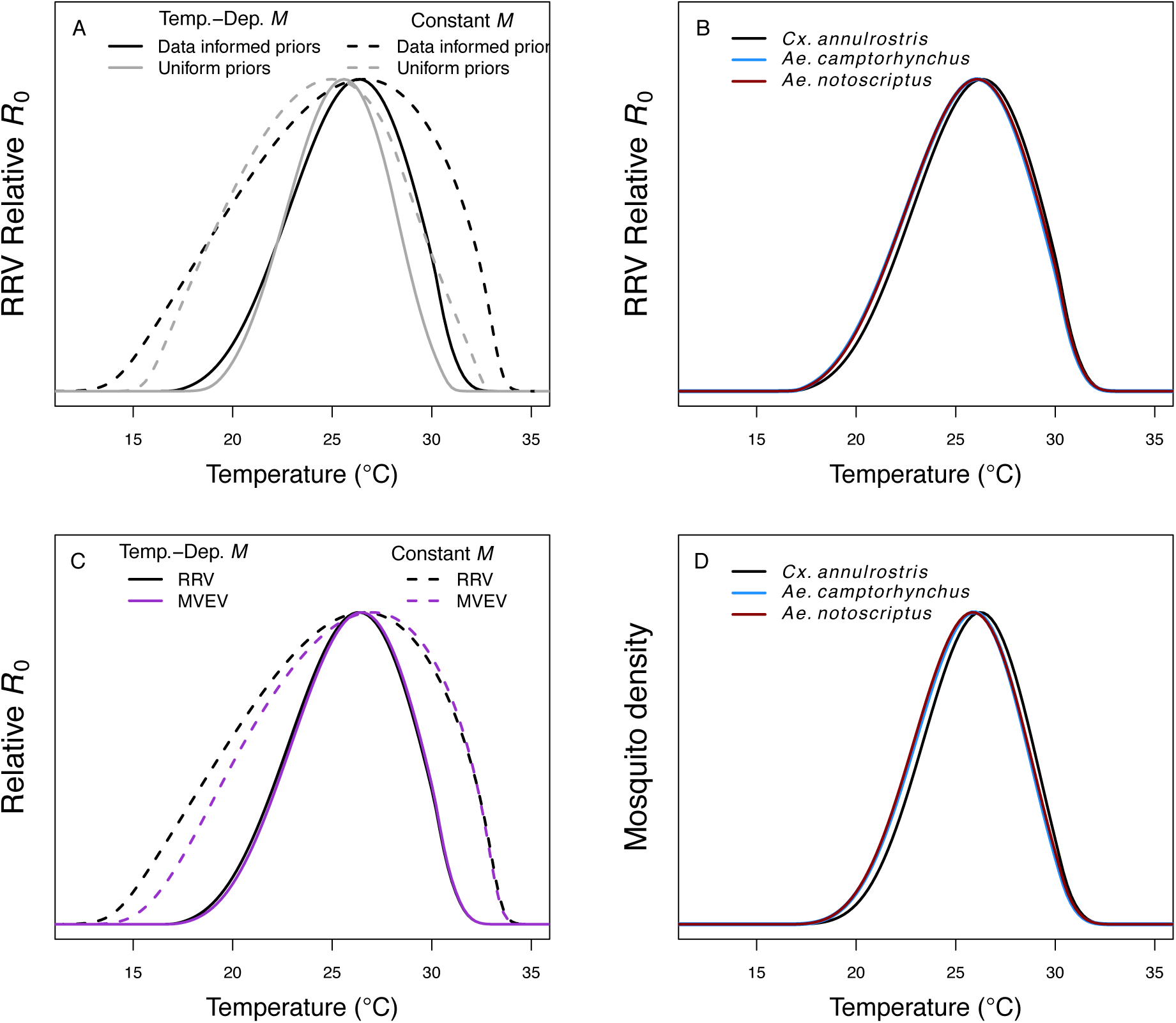
Additional *R_0_* and mosquito density model results. (A) *R_0_* (temperature-dependent *M* model, eq. 2: solid lines; constant *M* model, eq. 1: dashed lines) for Ross River virus (RRV) from traits fit with data-informed priors (black) and uniform priors (gray). (B) *R_0_* for RRV with partial trait data (*pLA* and *MDR*) from alternative vector species *Ae. camptorhynchus* (blue line) and *Ae. notoscriptus* (red line). Model using traits from *Cx. annulirostris* (solid black line) shown for comparison. These traits only occur in the temperature-dependent *M* model. (C) *R_0_* (temperature-dependent *M* model: solid lines; constant *M* model: dashed lines) for RRV (black) and Murray Valley Encephalitis virus (MVEV; purple). (D) Equilibrium mosquito density (*M*) from partial trait data (*pLA* and *MDR*) from alternative vector species *Ae. camptorhynchus* and *Ae. notoscriptus*. Lines are the same as in panel B.

**FIGURE S6:**
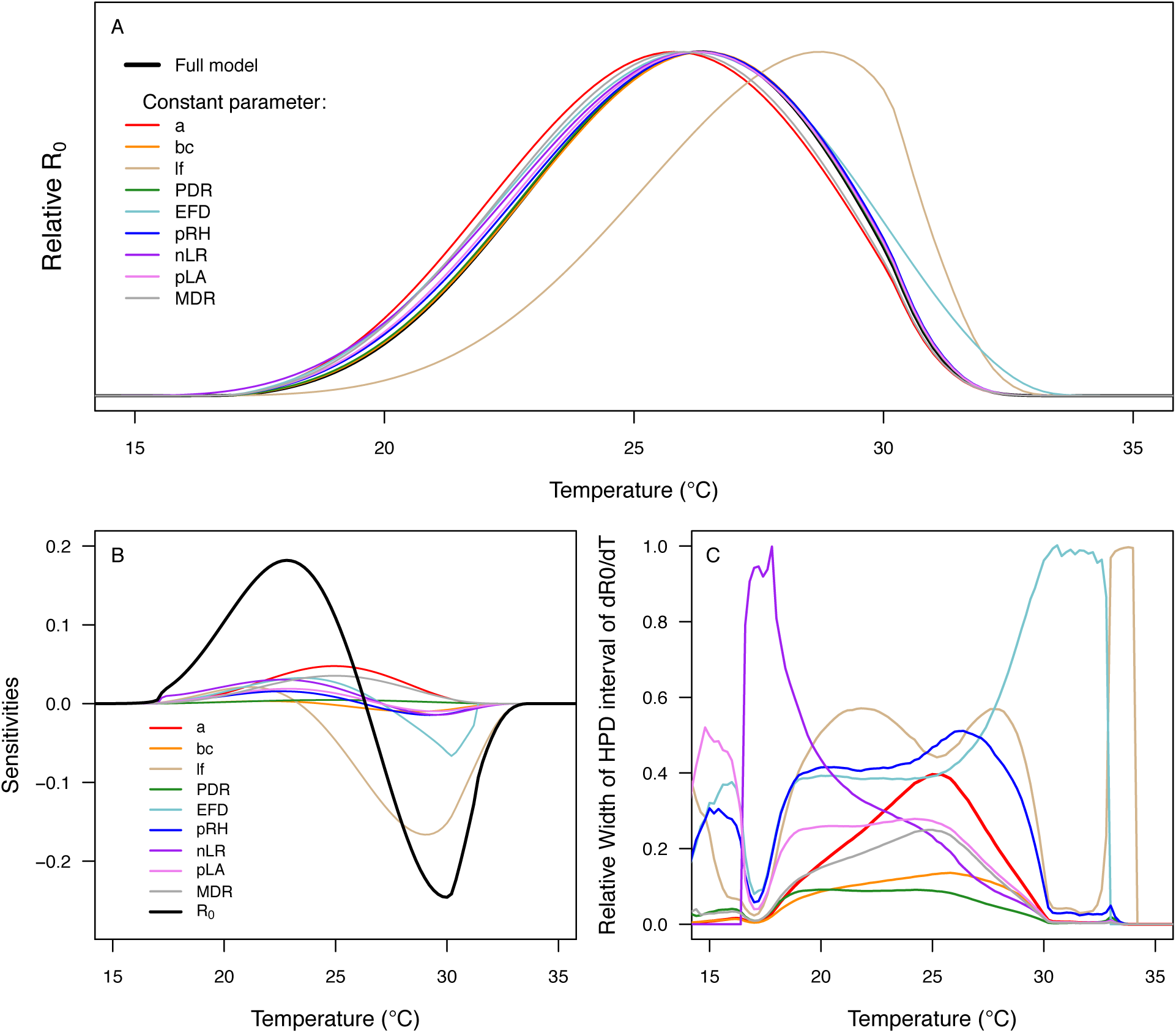
Sensitivity and uncertainty analyses of the temperature-dependent *M* model (eq. 2) to examine the effects of each trait’s thermal response on the thermal response of *R_0_*. (A) Sensitivity measured by holding one parameter constant while other parameters assume their posterior median thermal response functions. The black curve is the posterior mean of *R_0_*. Constant parameters are: biting rate (*a*, red), vector competence (*bc*, orange), fecundity (*EFD*, tan), proportion of egg rafts hatching (*pRH*, green), number of larvae emerging per raft (*nLR*, light blue), larval-to-adult survival (*pLA*, dark blue), mosquito development rate (*MDR*, purple), lifespan (*lf* = 1/*μ*, pink), and parasite development rate (*PDR*, gray). If setting a trait constant results in *R_0_* shifting to a higher optimum, then that trait is responsible for lowering the optimum of *R_0_* (and vice-versa). (B) Sensitivity measured as the partial derivatives of *R_0_* with respect to each trait and temperature. The thick black curve is the derivative of *R_0_* with respect to temperature; traits are as in panel A. (C) The relative contribution of each trait to uncertainty in *R_0_* across temperatures. (See Material and Methods in main text for more details.)

**FIGURE S7:**
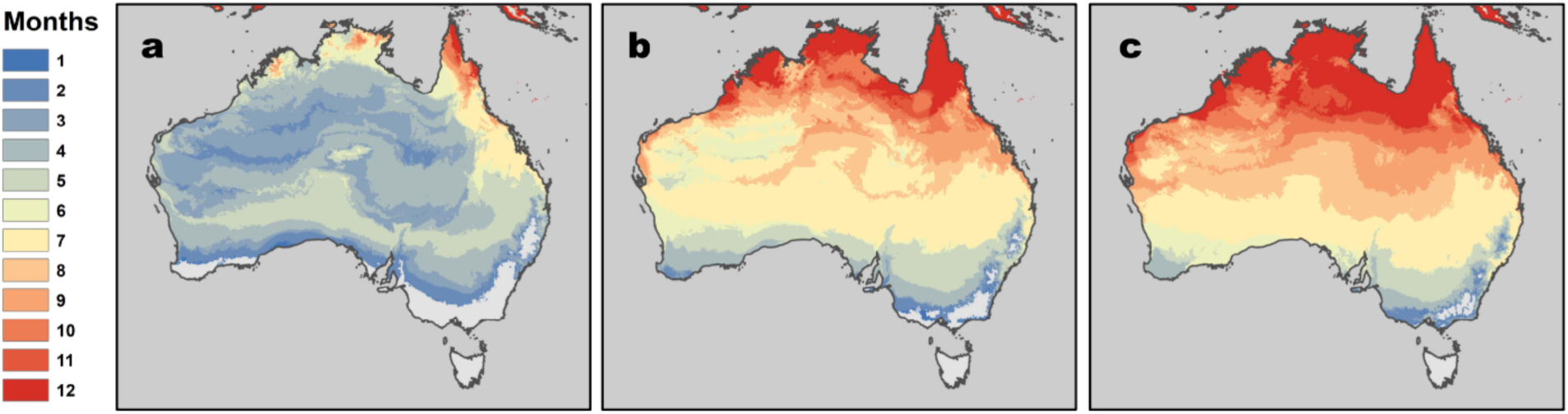
Maps showing CIs of temperature suitability for RRV transmission. Months where *R_0_* > 0 with (A) 2.5%, (B) 50% (median), and (C) 97.5% of the posterior probability distribution. Panel B is the same as Fig. 4A in the main text.

**FIGURE S8:**
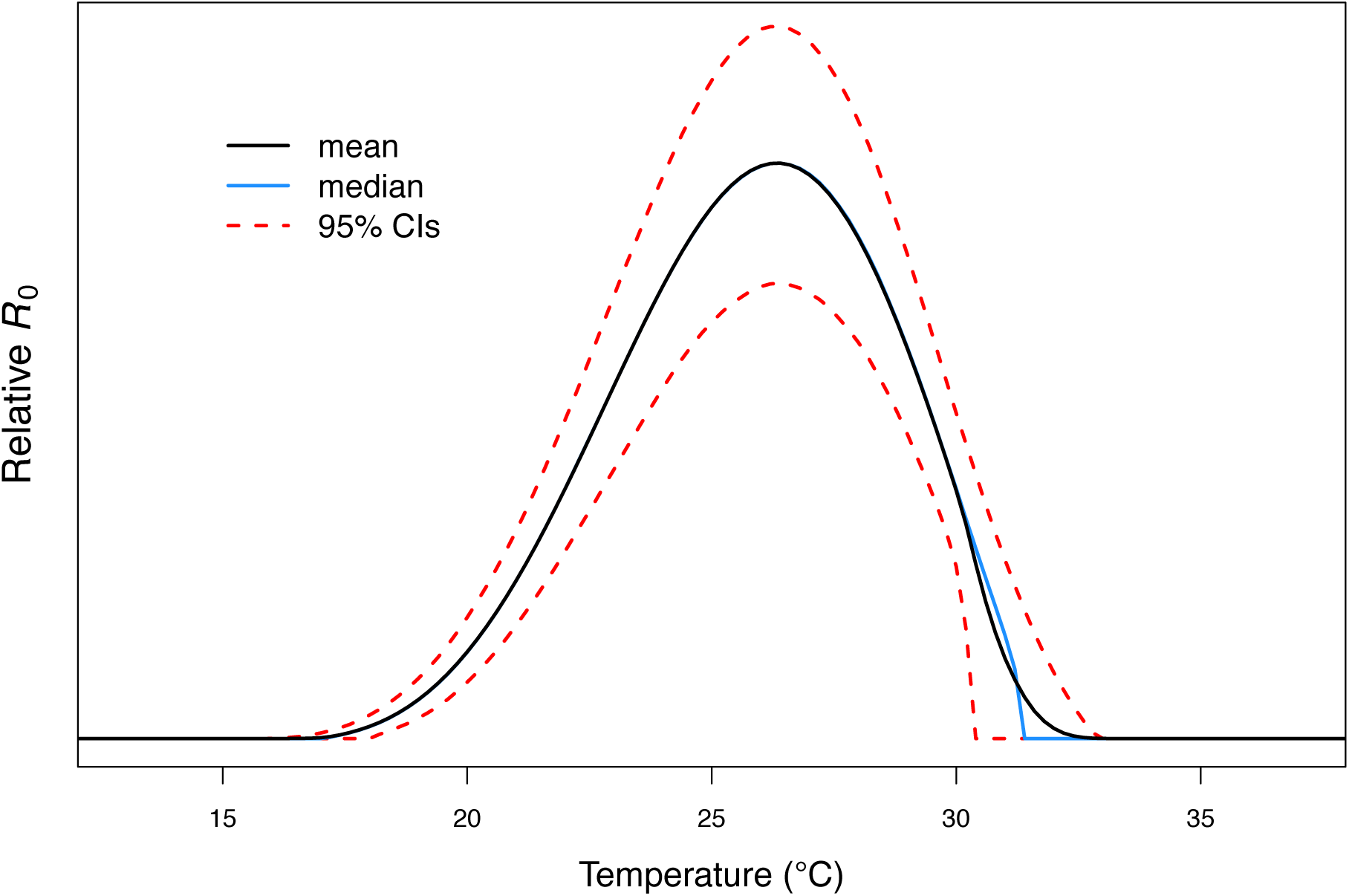
Mean versus median RRV *R_0_* results. Example showing how the upper thermal limit smoother and more visually intuitive with posterior distribution mean rather median. Throughout the main text and Appendix S1 we plot posterior distribution means but report medians of parameter values and use medians for downstream analyses (see note above).

## Partial Derivatives for Sensitivity Analysis

For all traits (*x*) that appear only in the numerator of *R_0_* (*bc, EFD, pRH, nLR, pLA, MDR*):

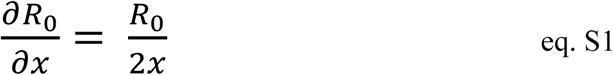

For biting rate (*a*):

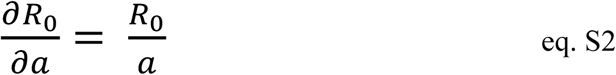

For parasite development rate (*PDR*):

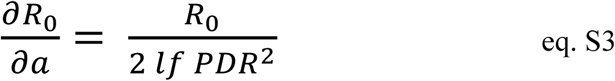

For lifespan (*lf*):

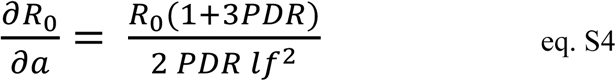

## Additional methods for survival and fecundity data from McDonald *et al*. 1980

The fecundity and adult survival data in McDonald *et al*. 1980 were both published as time series for one experimental population number over time. This approach resulted in data that needed to be formatted or transformed to better fit thermal responses via Bayesian inference.

For survival, McDonald *et al*. reported the percent surviving approximately every other day (hereafter: ‘semi-daily’). We used these data——along with the number of female adults alive on the first day of oviposition at each temperature—to generate a time series estimating the number surviving on every other day. To generate the dataset that we used to directly fit the thermal responses, we converted this time series into the number of female individuals who died on each day (i.e., lifespan data).

For fecundity, McDonald *et al*. reported semi-daily fecundity data for entire population. Because the population was synchronized, and because mosquitoes lay discrete clutches of eggs separated by several days (the gonotrophic cycle duration), there were many days in the time series in which the population did not produce offspring. These zero-inflated semi-daily fecundity data are not ideal for fitting thermal responses. Therefore, after digitizing the daily fecundity time series, we binned periods of several days (the bin size varied by temperature, since the gonotrophic cycle duration varies with temperature) and took a survival-weighted average within each bin (so days with more individual mothers contributing to offspring production counted more). To generate the dataset that we used to directly fit the thermal responses, we weighted the values within each bin by the mean number of surviving mothers in that bin. This approach allowed us to remove zero points and more accurately reflect daily fecundity averaged over a non-synchronized mosquito population. NOTE: the variation captured by these data and this approach is not variation between individual adult females, but rather variation by age for the entire population.

